# IL-9 as a naturally orthogonal cytokine with optimal JAK/STAT signaling for engineered T cell therapy

**DOI:** 10.1101/2025.01.15.633105

**Authors:** Hua Jiang, Sam Limsuwannarot, Kayla R. Kulhanek, Aastha Pal, Lea W. Rysavy, Leon Su, Ossama Labiad, Stefano Testa, Heather Ogana, Deepa Waghray, Pingdong Tao, Kevin M. Jude, Christopher S. Seet, Gay M. Crooks, Everett J. Moding, K. Christopher Garcia, Anusha Kalbasi

**Author notes:** These authors contributed equally to this work (Hua Jiang, Sam Limsuwannarot, Kayla R. Kulhanek, Aastha Pal, Lea W. Rysavy).

## Abstract

Arming T cells with a synthetically orthogonal IL-9 receptor (o9R) permits facile engraftment and potent anti-tumor functions. We considered whether the paucity of natural IL-9R expression could be exploited for T cell immunotherapy given that, in mice, high doses of IL-9 were well-tolerated without discernible immune modulation. Compared to o9R, T cells engineered with IL-9R exhibit superior tissue infiltration, stemness, and anti-tumor activity. These qualities are consistent with a stronger JAK/STAT signal, which in addition to STAT1/3/5, unexpectedly includes STAT4 (canonically associated with IL-12 but not common γ-chain cytokines). IL-9R T cells are exquisitely sensitive to perturbations of proximal signaling, including structure-guided attenuation, amplification, and rebalancing of JAK/STAT signals. Biased IL-9R mutants uncover STAT1 as a rheostat between proliferative stem-like and terminally differentiated effector states. In summary, we identify native IL-9/IL-9R as a natural cytokine-receptor pair with near-orthogonal qualities and an optimal JAK/STAT signaling profile for engineered T cell therapy.

## INTRODUCTION

Common γ-chain (γ_c_) cytokines are critical regulators of T cell differentiation and function, exerting their effects through JAK/STAT signaling pathways^1^. Their potential to enhance adoptive T cell therapy for cancer has long been recognized^2^, yet clinical translation of γ_c_ and other cytokines has been impeded by safety concerns stemming from their pleiotropic effects^3–7^. To overcome this challenge, several approaches have been developed to selectively deliver cytokine signals to engineered T cells. In one approach, mutations in IL-2 and its receptor, IL-2RB, yielded orthogonal IL-2 (oIL-2) and orthogonal IL-2RB (o2R), a synthetic and mutually exclusive pair^8^. This system enables targeted IL-2 signaling in engineered T cells while avoiding activation of other IL-2-responsive populations that contribute to systemic toxicity^9^.

To explore which of the γ_c_ cytokines induce the optimal JAK/STAT signals for successful T cell immunotherapy, we recently tested an iteration of the orthogonal IL-2 system in which we replaced the intracellular domain (ICD) of o2R with ICDs of different γ_c_ cytokine receptors (IL-4R, IL-7R, IL-9R, IL-21R)^10^. These chimeras (o4R, o7R, o9R, o21R) mimic the unique JAK/STAT signaling pattern of their natural receptor counterpart and induce different phenotypic and functional profiles in T cells. We observed that o9R, which induces a distinct JAK/STAT signaling profile consisting of STAT1, STAT3, and STAT5 activation, results in T cells that acquire stem-like characteristics, efficiently infiltrate tumors, and substantially improve the anti-tumor activity of TCR- and CAR-T cells in immunocompetent hosts even without conditioning lymphodepletion.

Given the relative anonymity of IL-9R signaling among the γ_c_ family, this surprising result prompted us to explore anti-tumor effects of T cells signaling through the wildtype IL-9 receptor. In particular, we were intrigued by the reported low nanomolar affinity of IL-9 for the wildtype IL-9 receptor^11^. However, it was not clear whether there is sufficient IL-9R expression on T cells to permit a response to IL-9. And furthermore, as the only γ_c_ cytokine not tested in humans, potential toxicities of systemic IL-9 administration are unknown.

To address these issues, we examined publicly available and in-house genomic, transcriptomic and functional data to characterize the expression and function of IL-9R in normal tissues. Compared to receptors for other common γc cytokines, IL-9R is not only expressed at very low levels in T cells, but across all tissues, including throughout fetal development, at different stages in the thymus, and in the tumor microenvironment. In fact, population-based genomic data suggests that IL-9 may be among the least understood cytokines from an evolutionary perspective.

Based on these and other findings, we nominated the wildtype IL-9 and IL-9 receptor (IL-9R) as a naturally orthogonal cytokine-receptor pair. Since IL-9 was well-tolerated in mice even at very high doses, we engineered T cells with the wildtype IL-9 receptor, and compared to T cells with o9R, found they have superior engraftment, tumor infiltration, stemness and anti-tumor effects in lymphoreplete mice without conditioning lymphodepletion. We linked these findings to more potent phosphorylation of STAT1, STAT3 and STAT5, as well as phosphorylation of STAT4, which is not classically associated with common γc signaling.

Through a multi-faceted approach to attenuate, amplify or rebalance the JAK/STAT signaling of IL-9R, we linked proximal signaling perturbations of IL-9R-engineeered T cells to their in vivo biologic outputs. Ultimately, the effects of IL-9R signaling on anti-tumor efficacy of T cells were exquisitely sensitive to both the strength and balance of JAK/STAT signaling. In particular, biased IL-9R mutants clarified the T cell intrinsic effect of STAT1 in tumor-specific T cells, skewing T cells from a stem-like state with proliferative capacity toward a terminally differentiated effector state. We further extend these results to human model systems using CAR T cells, setting the framework for a natural cytokine-receptor pair to mimic an orthogonal system and potentiate the anti-tumor efficacy of T cells in vivo.

## RESULTS

### The expression and activity of IL-9R is restricted in T cells and across normal tissues

To investigate the expression of IL-9R in normal T cells we examined single cell RNA-sequencing (scRNA-seq) of 28,964 peripheral blood mononuclear cells (PBMCs) from two healthy donors, capturing major hematopoietic lineage cell types (Figure 1A)^12^. Compared to receptors for other common γc cytokines IL-2/IL-15 (*IL2RB*), IL-4 (*IL4R*), IL-7 (*IL7R*), and IL-21 (*IL21R*), *IL9R* was expressed at low levels and in very few cells. *IL9R* transcripts were detected in rare B cells (0.53%) and CD4 T cells (0.26%). We expanded our search to tissue resident immune cells using a scRNA-seq dataset of 39,560 lamina propria immunocytes (Figure 1B)^13^. Consistently, *IL9R* was expressed rarely and at low levels, in contrast to *IL2RB, IL4R, IL7R,* and *IL21R*.

**Figure 1.**
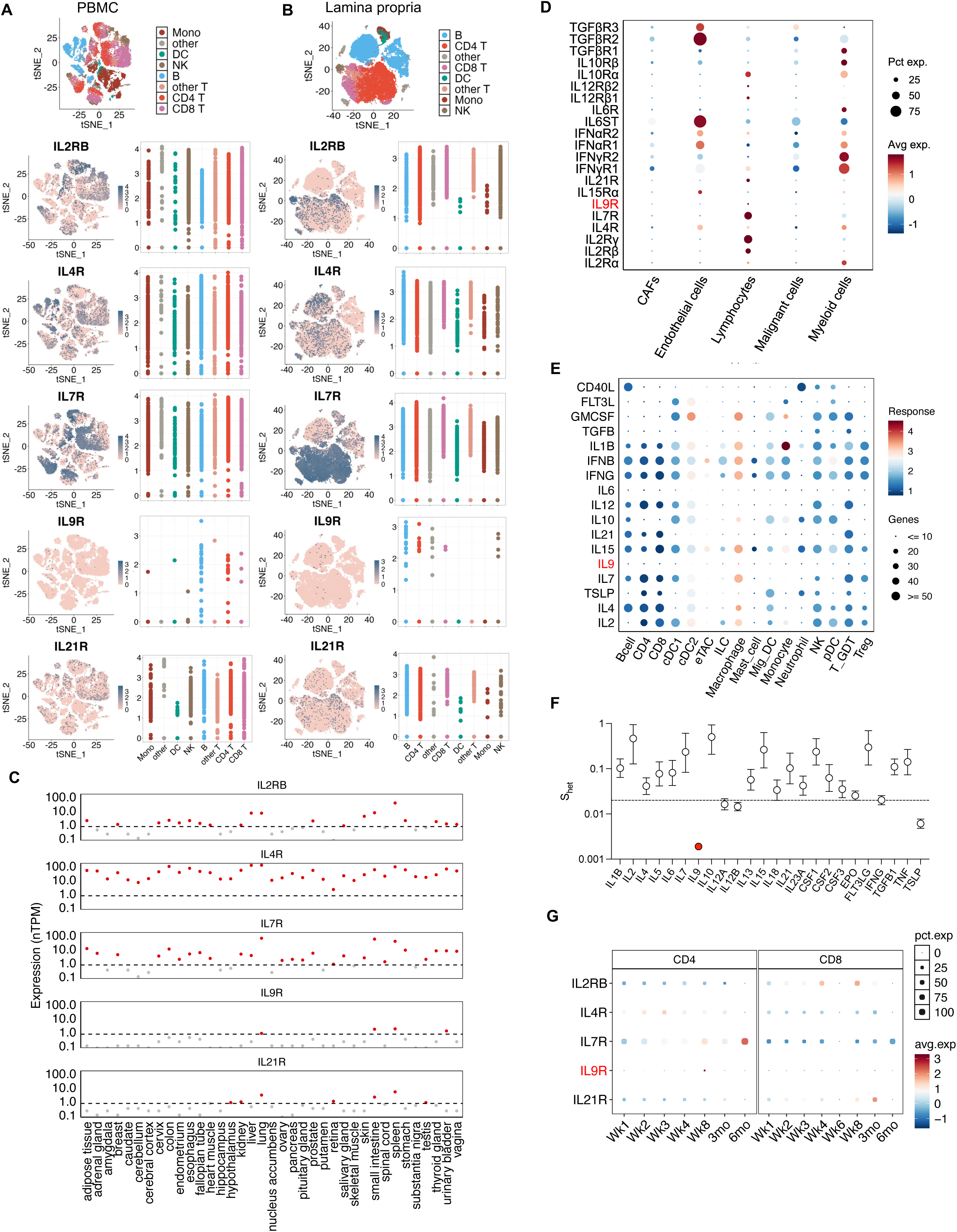
Nomination of IL-9 as a naturally orthogonal cytokine for T cell therapy. (A) Expression of common γ_c_ cytokines receptors in single cell RNA sequencing data of PBMC from normal healthy donors. tSNE plots with cell annotations are shown in the top panel as a reference. For each cytokine receptor, a feature plot colored by scaled expression level (left panel) and a scatter plot summarizing expression for each cell type (right panel) are shown. (B) As in (A), but for lamina propria lymphocytes from the uninflamed ileum of patients with Crohn’s disease. (C) Expression scatterplot of normalized expression of receptors for γ_c_ cytokines in publicly available RNA sequencing data of 37 normal tissues from the GTEx database. Dashed line indicates a strict nTPM threshold of 0.7, below which expression is not detected or negligible. (D) Dot plot of intratumoral expression of a panel of cytokine receptors across malignant and non-malignant cell types, based on single cell RNA-sequencing data from soft tissue sarcoma (n=65,945 cells across 15 patients). (E) Differential expression of genes (number of genes, magnitude of response) among cell types within lymph nodes of mice treated with one of several cytokines versus PBS control (based on publicly available data). (F) Reported median S_het_ values for γ_c_ and other cytokines. S_het_ quantifies the impact of predicted heterozygous loss-of-function variants on evolutionary fitness, with higher scores associated with greater impact on evolutionary fitness. Error bars represent 95% highest posterior density. The dashed line at 0.02 represents the median S_het_ value for all canonical transcripts. (G) Expression dot plot of receptors for γ_c_ cytokines in sorted CD19 CAR T cells (n=66,042) from 15 pediatric patients at various timepoints after infusion. Data presented separately for CD4 (left) and CD8 (right) CAR T cells. PBMC: peripheral blood mononuclear cells; TPM: transcripts per million; GTEX: Genotype-Tissue Expression; CAR: chimeric antigen receptor

Even throughout stages of thymic development, *IL9R* transcript levels are nearly absent (Figure S1A)^14^. Consistent with the low transcript expression in the vast majority of developing and mature T cells, activated human T cells from three independent healthy donors did not phosphorylate STAT proteins in response to IL-9 (Figure S1B).

We extended the analysis beyond immune cells to 37 human tissue types using publicly available bulk RNA-sequencing data from the Genotype-Tissue Expression (GTEx) project^15,16^. Normal tissue expression of *IL9R* is broadly restricted, with only four of 37 tissues having *IL9R* expression above the threshold of log_10_(nTPM)>0.7: lung, small intestine, spleen and urinary bladder (Fig. 1C). Restricted *IL9R* expression was also evident during fetal development (Figure S1C).

It is plausible that inflammatory cues, such as those within the tumor microenvironment, induce *IL9R* expression. However, using scRNA-seq data from soft tissue sarcoma samples (n=65,945 cells across 15 patients), we also observed restricted expression of *IL9R* across major cell types in the tumor microenvironment, compared to a panel of 20 other canonical cytokine receptors (Fig. 1D).

We next sought functional readouts to assess the potential toxicity of systemically administered IL-9. For this, we evaluated publicly available scRNA-seq data obtained from lymph nodes of mice treated with one of 86 different cytokines (n=3 mice per cytokine) or PBS control (n=14 mice)^17^. For each cell type, we evaluated differential gene expression in response to each cytokine compared to PBS control treated mice to ascertain the magnitude of the global transcriptomic response as well as the total number of significantly upregulated genes. In contrast to both γc and non-γc cytokines, IL-9 resulted in minimal impact on gene expression profiles compared to control treated mice, with no more than one differentially expressed gene (absolute log_2_(fold change)>0.25, FDR-adjusted *P*<0.05, two-sided Wilcoxon rank-sum text) in any cell cluster (Fig. 1E).

We hypothesized that IL-9 may have a lesser role in evolutionary fitness compared to other cytokines. To explore this, we utilized analysis from population-scale exome sequencing data (n=983,578 individuals) from the Regeneron Genetics Center Million Exome (RGC-ME) project. This analysis reports a coefficient, s_het_, using predicted heterozygous loss-of-function mutations, which captures the evolutionary indispensability of protein-coding genes^18,19^. IL-2, IL-7 and IL-15 were among cytokines with the highest estimated indispensability (S_het_). In contrast, among a panel of canonical cytokines, IL-9 ranked as the most dispensable with a mean s_het_ value of 0.0019 (95% highest posterior density: [0.0017, 0.0022]), well below the mean for all canonical transcripts (0.073)(Fig. 1F).

Given the absence of sufficient IL-9R expression on T cells, exploiting the IL-9/IL-9R axis for T cell therapy would require genetic modification of T cells. To confirm that current manufacturing strategies do not induce IL-9R expression, we examined scRNA-seq from 66,042 post-infusion CAR T cells obtained from 16 pediatric patients at various time points after CAR T cell administration^20^. Unlike the expression of *IL2RB and IL21R* (primarily observed in CD8+ CD19 CAR T cells) or *IL4R* and *IL7R* (primarily observed in CD4+ CD19 CAR T cells), *IL9R* expression was notably absent from CD19 CAR T cells (Fig. 1G).

### IL-9 is well-tolerated and drives potent anti-tumor efficacy of T cells engineered with IL-9R

To evaluate the in vivo effects of IL-9, we engineered a half-life extended version of mouse IL-9 by tagging mouse serum albumin to the N-terminus (MSA-IL9), which, in contrast to MSA-IL2, was well-tolerated by metrics of weight loss, survival and mobility, even at an excess dose of 100μg given every other day for three weeks (Fig. 2A-D). Whereas MSA-IL2 increased serum IFNγ levels, MSA-IL9 did not (Fig. 2E).

**Figure 2.**
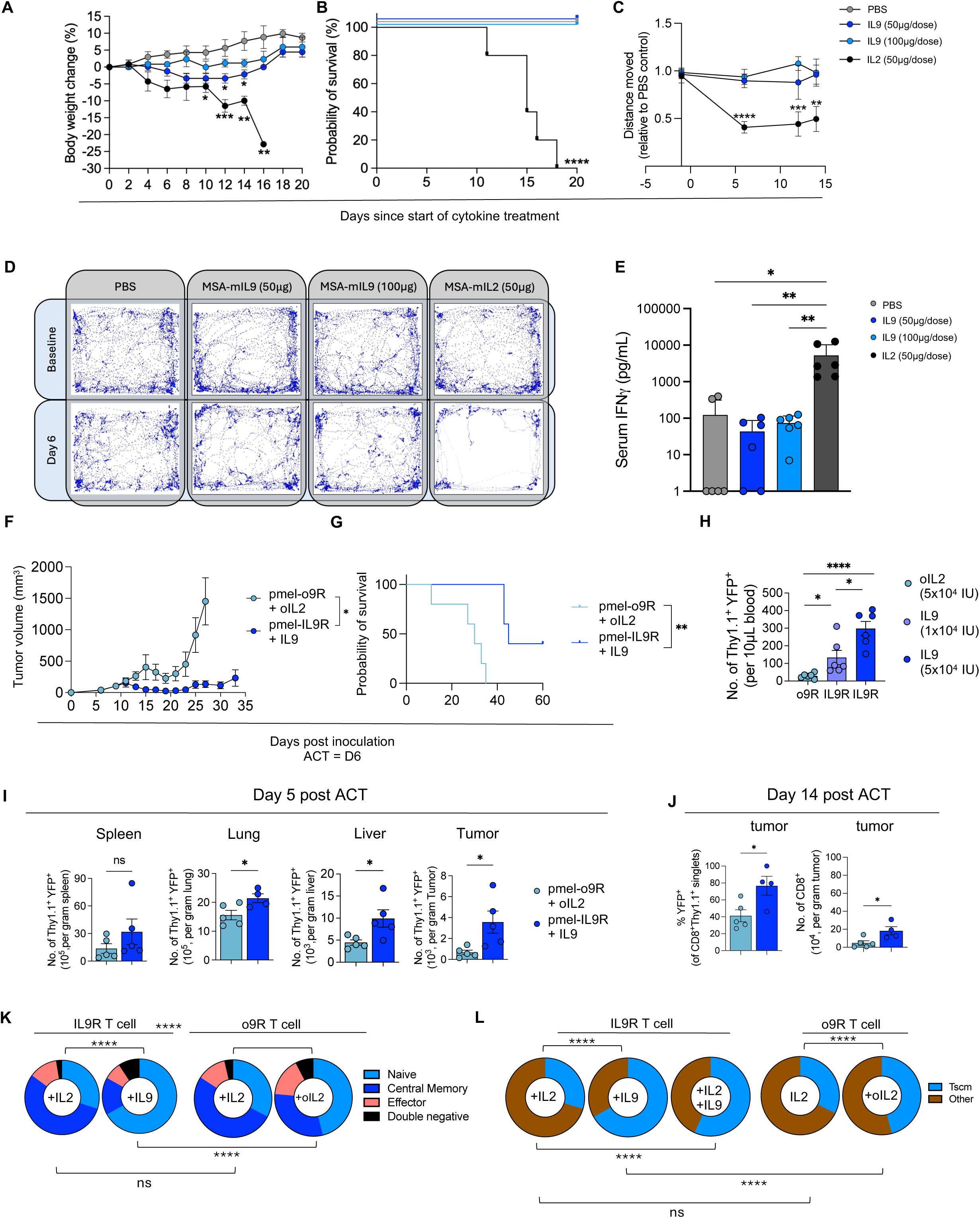
Exogenous IL-9 is well-tolerated and results in superior tissue infiltration, stemness and anti-tumor activity of IL-9R-engineered antigen-specific T cells. (A) Body weight (percentage change from baseline) of mice (n=8 mice/group) treated with PBS, IL-9 (50 µg or 100 µg), or IL-2 (50 µg) i.p. every other day. (B) Survival of mice from (A). (C) Sum of distance traveled over 30 second period in mice from (A), plotted relative to the average distance traveled by mice in the PBS group. (D) Graphical representation of mouse movement (n=8 mice/group) over a period of 30 seconds at baseline (top panel) and six days after starting treatment (bottom panel). (E) Serum IFNγ levels measured by ELISA six days after initiating treatment (n=3 mice/group) as described in A-C. (F) B16-F10 tumor growth (mean ± SEM, n=5 mice/group) after ACT with pmel T cells engineered with o9R or IL-9R (0.4×10^6^ transduced cells, i.v.), and cytokine treatment with oIL-2 or IL-9, respectively (5×10^4^ IU i.p., daily for 5 days starting with ACT). Data is representative of three independent experiments. (G) Survival of mice from (F). (H) Quantification of transduced pmel T cells in the blood five days after ACT (n=5 mice/group). (I) Quantification of transduced pmel T cells (Thy1.1^+^YFP^+^) across different tissues five days after ACT (n=5 mice/group). (J) Transduced (YFP^+^) pmel T cells as percentage of all pmel T cells within the tumor 14 days after ACT (left panel) and total number of CD8+ T cells per gram of tumor 14 days after ACT (right panel) (n=5 mice per group). (K) o9R or IL9R transduced C57BL/6 T cells stimulated for 24h with IL-2 (10nM), IL-9 (10nM), oIL-2 (10µM), or a combination as noted. The proportion of naïve (CD62L+CD44-), central memory (CD62L+CD44+), effector (CD44+CD62L-) or double negative (CD62L-CD44-) T cells. Data from two independent experiments, with 4 technical replicates per condition. (L) The proportion of T_SCM_ cells (CD44-, CD62L+, Sca-1+) from (K). For in vivo experiments, cytokines were tagged with mouse serum albumin (MSA) for half-life extension. *P < 0.05, **P < 0.005, ***P < 0.0005, ****P < 0.0001 (one-way ANOVA for A-C and E-F; Mantel-Cox for G; unpaired t test for H and J, two-way ANOVA for K-L).

We then compared the anti-tumor efficacy of H2-D^b^/gp100-specific T cells from pmel-1 transgenic mice (hereafter, pmel mice) engineered with either IL-9R or our benchmark, the synthetic o9R, in B16-F10 tumor-bearing mice treated with the corresponding cytokine (MSA-IL9 or MSA-oIL2). IL-9R pmel T cells resulted in superior anti-tumor efficacy and prolonged survival compared to o9R pmel T cells (Fig. 2F-G, Fig. S2A). We attributed this primarily to the effects of MSA-IL9 on the IL-9R pmel T cells, as MSA-IL9 alone did not result in tumor growth delay (Fig. S2B). We validated the anti-tumor potency of IL-9R pmel T cells in a mouse model of sarcoma derived from autochthonous tumors in genetically modified Kras^G12D/-^P53^flox^ mice (Fig. S2C).

The superior anti-tumor efficacy of IL-9R pmel T cells was associated with improved peripheral expansion compared to o9R pmel T cells. Peripheral expansion was IL-9 dose-dependent, supporting the cytokine-dependent activity of the IL-9R pmel T cells (Fig. 2H). Beyond peripheral expansion, we also noted enhanced early tissue infiltration by IL-9R pmel T cells compared to o9R pmel T cells, including the lung, liver and tumor (Fig. 2I). And although IL-9R pmel T cells were not enriched compared to o9R T cells in normal tissues after cytokine withdrawal (Fig. S2D), IL-9R pmel T cells continued to be enriched at the tumor site (Fig. 2J).

We previously demonstrated^10^ that o9R signaling selectively expands T cells with a stem cell memory (T_SCM_) phenotype. Here, we observed that IL-9R signaling resulted in more efficient induction of naïve CD62L^+^CD44^-^ T cells (Fig. 2K, Fig. S2E-F), including the T_SCM_ (CD62L^+^CD44^-^ Sca-1^+^) subset (Fig. 2L, Fig. S2G), even in the presence of IL-2. Together, these results indicate that compared to the synthetic o9R system, natural IL-9R signaling further promotes stemness and tumor infiltration, and results in more potent anti-tumor functions of T cells.

### In addition to robust activation of STAT1, STAT3 and STAT5, IL-9R recruits STAT4 signaling

We sought to elucidate the unique signaling characteristics of IL-9R that contribute to its enhanced anti-tumor activity, first comparing the signaling strength of IL-9R and o9R. Dose-response curves of STAT1, STAT3, and STAT5 phosphorylation revealed that IL-9R resulted in higher E_max_ and lower EC_50_ than o9R, indicating both higher affinity and stronger STAT signaling (Fig. 3A). These differences in signaling strength resulted in differential transcriptomic outputs based on RNA-sequencing 48h after cytokine stimulation in vitro. IL-9R T cells treated with IL-9 clustered separately from o9R T cells treated with oIL-2 on principal component analysis (Fig. S3A). However, in contrast to the transcriptomic output of IL-2 signaling, the transcriptomic outputs of IL-9R and o9R were relatively similar (Fig. 3B, Fig. S3B). While 7673 differentially expressed genes (DEGs) were observed between IL-9R + IL-9 and IL-9R + IL-2 groups, and 7166 DEGs between o9R + oIL-2 and IL-9R + IL-2 groups, only 3031 DEGs were noted between IL-9R + IL-9 and o9R + oIL-2 groups (Fig. 3B, Fig. S3A-C). Thus, IL-9R and o9R activate similar gene expression patterns, consistent with their shared intracellular signaling domains, yet their transcriptomic profiles are distinguishable due to differences in proximal signal strength.

**Figure 3.**
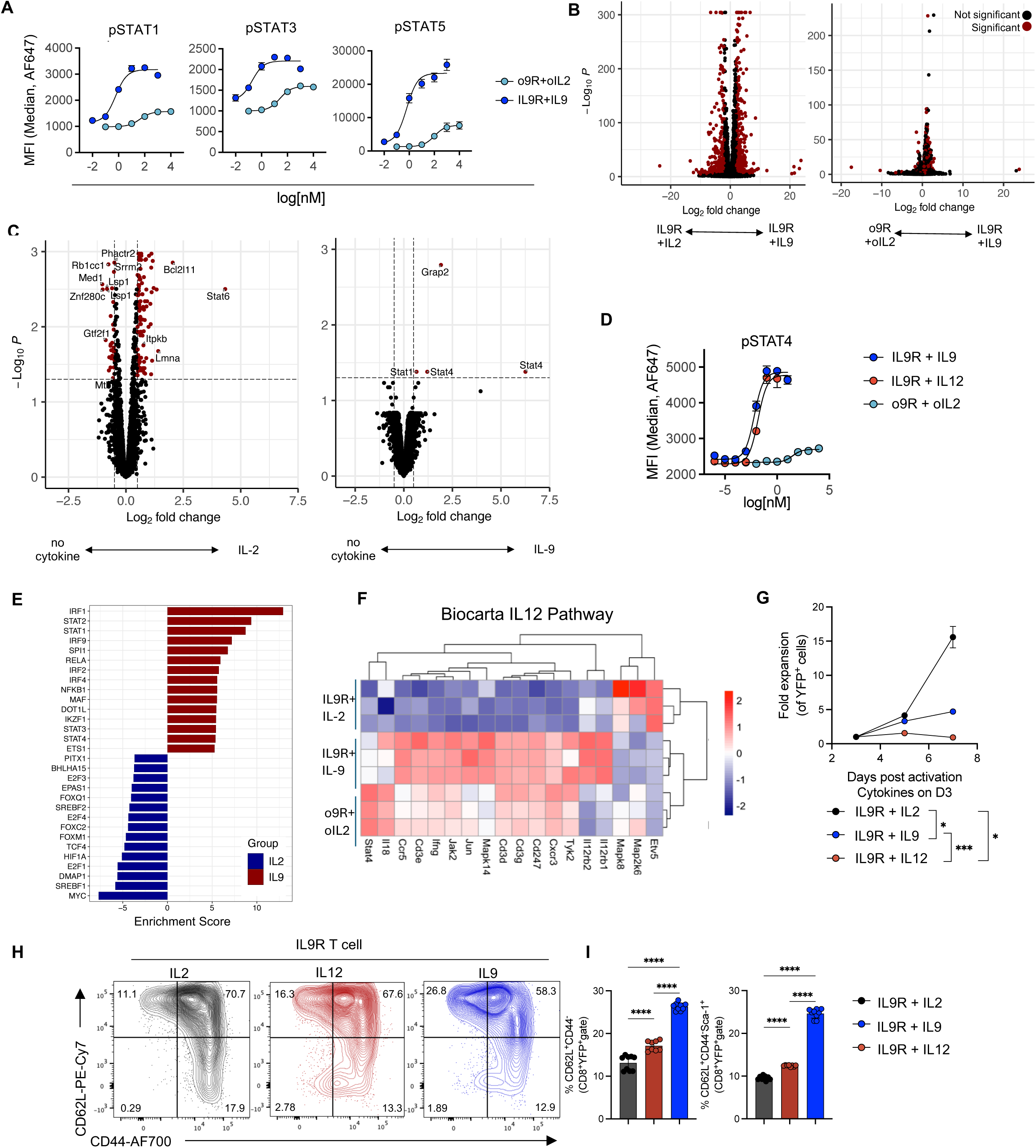
IL-9R signaling results in phosphorylation of STAT4, in addition to STAT1, STAT3, and STAT5. (A) Dose-response curves of STAT1, STAT3, and STAT5 phosphorylation in IL-9R or o9R transduced (YFP^+^) pmel T cells stimulated with either oIL-2 or IL-9 for 20 minutes (shown are technical duplicates; representative of at least three independent experiments). (B) Volcano plots depicting differential gene expression based on RNA-sequencing of IL-9R or o9R transduced C57BL/6 T cells and treated with IL-9 (10 nM), IL-2 (10 nM) or oIL2 (10 µM) for 48 hours. Comparisons for each volcano plot are shown below the x-axis. Significance (red) indicates adjusted p < 10^-5^ and absolute fold change ≥2. (C) Differentially phosphorylated proteins between IL-9R transduced C57BL/6 T cells stimulated for 20’ with either IL-2 (10nM) or no cytokine (left) or IL-9 (10nM) versus no cytokine (right). Significance (red) indicates adjusted p<0.05 and log_2_(fold change)≥ 0.5. (D) Dose-response curve for STAT4 phosphorylation among IL-9R transduced (YFP^+^) pmel T cells treated for 20 minutes with IL-9 or IL-12, or o9R transduced T cells treated with oIL-2 (shown are technical duplicates; representative of two independent experiments). (E) Waterfall plot summarizing transcription factor enrichment scores based on RNA-seq data of IL-9R T cells treated with IL-9 or IL-2 (top 15 for each). Enrichment score is inferred by fitting a linear model that predicts observed gene expression based on prior knowledge of a curated set of transcription factors and their target genes. (F) Heat map of the expression of the Biocarta IL-12 Pathway gene set based on RNA-sequencing from (B). Samples and genes clustered hierarchically without supervision. (G) In vitro expansion of IL-9R (YFP^+^) transduced pmel T cells and treated with 10nM cytokine starting on day 3 after activation (n=3 technical replicates/group). Data is representative of three biological replicates and two independent experiments. (H) Representative contour plots of CD44 and CD62L expression of T cells from (G) after 24h treatment with cytokine. (I) Quantification of naïve CD62L^+^CD44^-^ (left) and stem-like CD62L^+^CD44^-^Sca-1^+^ (right) T cells from (H). Data from two independent experiments. *P < 0.05, **P < 0.005, ***P < 0.001, ****P < 0.0001 (one-way ANOVA for G; two-way ANOVA for G, I).

To explore properties of IL-9R beyond its known signaling through STAT1, STAT3 and STAT5, we performed phoshphoproteomic analysis of IL-9R-transduced T cells from C57BL/6 mice stimulated for 20 minutes with IL-9, IL-2, or no cytokine. Treatment with IL-2 resulted in a broad phosphoproteomic footprint, with 228 differentially abundant phosphoproteins, consistent with its known activation of multiple signaling pathways including JAK/STAT, ERK and AKT^21,22^ (Fig. 3C, Fig. S3D, Table S1). However, the phosphoproteomic effect of IL-9 was highly restricted, with only four differentially abundant phosphoproteins, with the JAK/STAT pathway accounting for three of the four differentially abundant phosphoproteins. With phosphoflow cytometry, we confirmed that unlike IL-2, IL-9 does not induce phosphorylation of ERK or AKT in IL-9R engineered T cells (Fig. S3E-F).

Unexpectedly, the most differentially abundant phosphoprotein upon IL-9 stimulation was pSTAT4, which has not been canonically implicated in IL-9 or, more broadly, common γ_c_ signaling (Fig. 3C)^1^. We validated this by phosphoflow, demonstrating comparable pSTAT4 activity to IL-12 signaling, the canonical activator of STAT4 in T cells^23^ and a well-described promoter of anti-tumor functions in T cells (Fig. 3D)^24–26^. Transcription factor enrichment analysis based on RNA-seq data further supported the role of STAT4 in IL-9R signaling, showing enrichment of STAT1, STAT2, STAT3, and STAT4 compared to IL-2 signaling (Fig. 3E). Additionally, both IL-9R and o9R signaling upregulate RNA expression of 15 of 18 genes in the Biocarta IL-12 Pathway gene set compared to IL-2 signaling (Fig. 3F).

While IL-12 exclusively signals through STAT4, IL-9 simultaneously activates STAT1, STAT3, STAT4 and STAT5. This distinction is evident in the functional outcomes of IL-9 and IL-12 signaling. In vitro expansion assays demonstrated that IL-9 provides a modest proliferative signal to IL-9R transduced T cells, unlike IL-12, but less potent than IL-2 (Fig. 3G), likely attributable to a higher STAT5 signal from IL-2^27^. Analysis of CD44 and CD62L expression revealed that IL-9 signaling results in enrichment of naïve T cells compared to IL-12, as well as enrichment of CD62L^+^CD44^-^Sca-1^+^ T_SCM_ cells (Fig. 3H-I), which has been associated with STAT3 signaling^28^.

In summary, these results demonstrate that IL-9R signaling recruits STAT4 in addition to robustly activating STAT1, STAT3, and STAT5. This unique combination of STAT activation leads to a distinct transcriptional profile and T cell phenotype that integrates features of IL-12 and γ_c_ cytokines, contributing to the anti-tumor efficacy observed with IL-9R-engineered T cells.

### Either attenuation or amplification of IL-9R signaling disrupt its anti-tumor properties

Given its unique JAK/STAT signaling profile, we sought to explore the effects of perturbations in proximal signaling on anti-tumor efficacy of IL-9R T cells. In other γ_c_ cytokines, a mutation in the glutamine residue in the helix D that interacts with the γ_c_ has been shown to attenuate signaling^29^. We thus selected to mutate the corresponding glutamine residue at position 115 of IL-9 to threonine (Q115T) (Fig. S4A). A predicted structure of the IL-9R complex also supported the interaction of Q115 with the γ_c_ (Fig. 4A). Indeed, we observed attenuated E_max_ and EC_50_ of STAT1, STAT3, STAT4, and STAT5 signaling of IL-9R engineered T cells in response to IL-9^Q115T^ compared to IL-9^WT^ (Fig. 4B).

**Figure 4.**
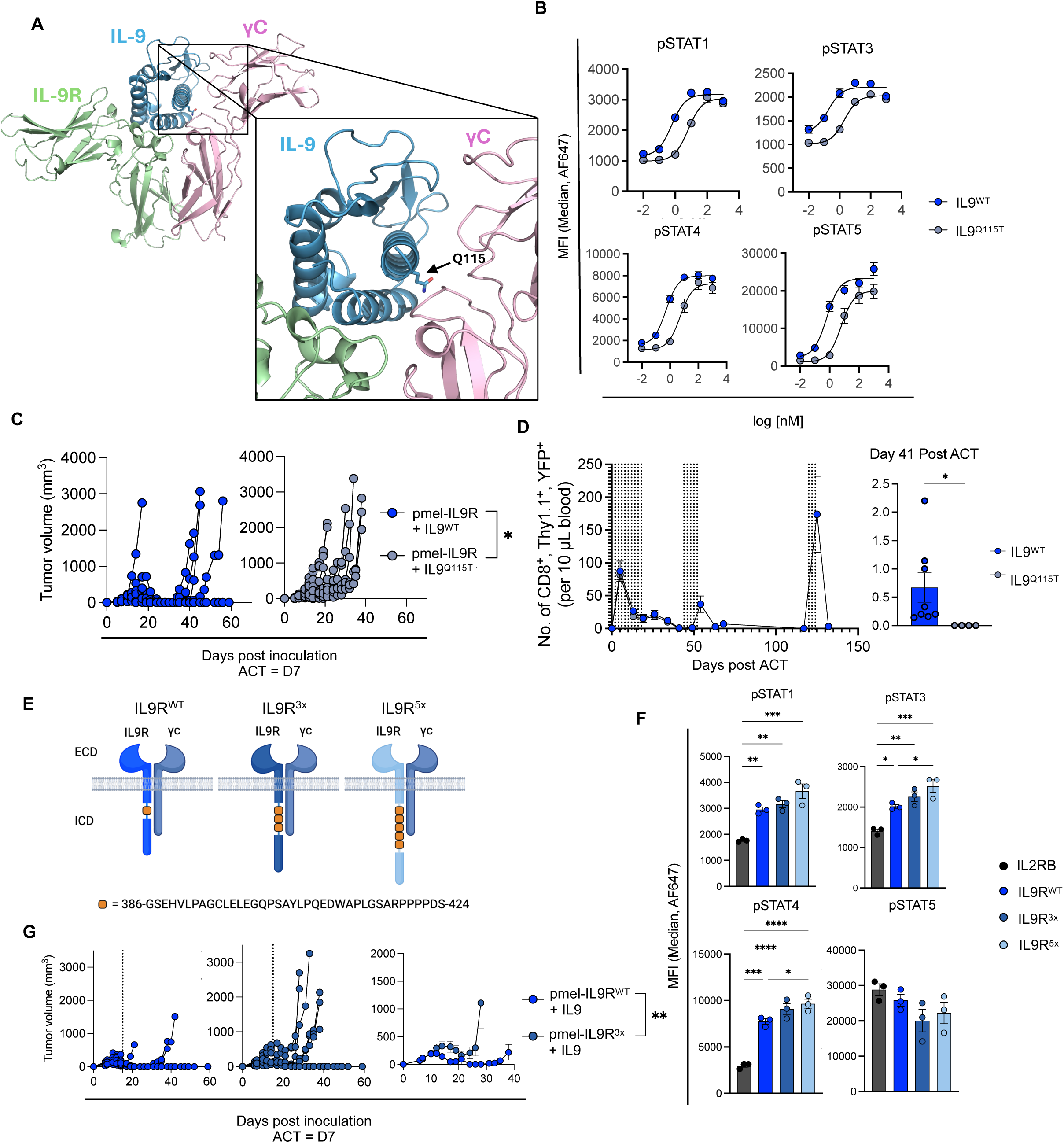
Structure-based attenuation or amplification of IL-9/IL-9R signaling diminishes the anti-tumor qualities of T cells signaling through the native receptor complex. (A) Structural prediction of the interleukin-9 (IL-9) receptor complex based on AlphaFold2. The complex consists of the IL-9 receptor (IL-9R, green), IL-9 (blue), and the γ_c_ (pink). The inset demonstrates the interaction of the glutamine at amino acid position 115 of IL-9 (Q115) with the γ_c_, which we subsequently mutated (IL-9^Q115T^) to generate an attenuated cytokine. (B) Dose-response curves of STAT1, STAT3, STAT4, and STAT5 phosphorylation in IL-9R transduced pmel T cells stimulated with IL-9^WT^ (blue) or IL-9^Q115T^ (light gray) for 20 minutes. Error bars represent SEM of technical duplicates; Data is representative of two independent experiments. (C) Tumor growth after ACT with IL-9R pmel T cells seven days after B16-F10 tumor inoculation. Mice were treated with either IL-9^WT^ (n=7 mice) or IL-9^Q115T^ (n=9 mice) (10 doses, every other day). Data is representative of two independent experiments. (D) Peripheral blood quantification of IL-9R transduced pmel T cells over time in mice treated with ACT and either IL-9^WT^ or IL-9^Q115T^ cytokine (each cytokine dose indicated by a vertical dashed line). In the IL-9^Q115T^ group, IL-9R pmel T cells were not observed in the blood at day 41 or after three additional cytokine doses on days 44-48, and thus only the IL-9^WT^ group received doses beyond day 48. Data is representative of two independent experiments. (E) Schematic of IL-9R variants with either three (IL-9R^3x^) or five (IL-9R^5x^) repeated phospho-tyrosine (pY) elements within the intracellular domain of the IL-9 receptor (created with Biorender.com). Sequence of the phosphotyrosine element shown in the legend. (F) Phosphorylation of indicated STATs at E_max_ (100nM) for IL-9R^WT^, IL-9R^3x^, and IL-9R^5x^ transduced pmel T cells stimulated with IL-2 or IL-9 for 20 minutes. Data is representative of three biological experiments (mean ± SEM). (G) B16-F10 tumor growth after ACT with either IL-9R^WT^ (n=7 mice) or IL-9R^3x^ transduced pmel T cells (n=8 mice). Mice were also treated with IL-9 (5×10^4^ IU i.p., every other day for 5 doses starting with ACT). Shown are individual tumor growth curves (left and middle) and mean ± SEM (right). Data is representative of two independent experiments. For in vivo experiments, cytokines were tagged with mouse serum albumin (MSA) for half-life extension. *P < 0.05, **P < 0.005, ***P < 0.001, ****P < 0.0001. (two-way ANOVA for C; Welch’s t-test for D, G; one-way ANOVA for F). See also Figure S4.

The subtle attenuation of IL-9 diminished the anti-tumor efficacy of IL-9R pmel T cells against B16 melanoma, both as measured by tumor growth and maximal treatment response (Fig. 4C, Fig. S4B). Although the initial expansion and persistence of IL-9R pmel T cells in the blood was similar in mice treated with IL-9^WT^ and IL-9^Q115T^, by day 41 we no longer observed IL-9R pmel T cells in mice treated with attenuated IL-9^Q115T^, whereas IL-9R pmel T cells in mice treated with IL-9^WT^ continued to persist (Fig. 4D, Fig. S4C). Furthermore, retreatment of mice with a second course of IL-9^WT^ resulted in a second peak of IL-9R pmel T cells in the blood, whereas retreatment with IL-9^Q115T^ did not. In fact, IL-9R pmel T cells were detectable over 115 days after initial adoptive transfer, at which point a third course of IL-9^WT^ resulted in a third peak of IL-9R pmel T cells, exceeding levels of the initial peak expansion.

To explore the effect of amplifying IL-9R signaling, we leveraged prior evidence that IL-9R signaling is driven by a single phosphotyrosine site (Y405) downstream of the Box 1 motif^30^. We generated IL-9R mutants in which a 39 amino acid sequence containing sequences flanking the phosphotyrosine site were repeated either three or five times (IL-9R^3x^ and IL-9R^5x^, respectively) (Fig. 4E). The mutant IL-9R^3x^ and IL-9R^5x^ increased the E_max_ of IL-9 for pSTAT1, pSTAT3, and pSTAT4 compared to IL-9R^WT^, though not STAT5 (Fig. 4F); these differences in pSTAT1, pSTAT3 and pSTAT4 reached significance for IL-9R^5x^ versus IL-9R^WT^ (Fig. S4D). As expected, the IL-9R^3x^ and IL-9R^5x^ did not impact the EC_50_.

Despite the subtle amplification of the IL-9R signal, the anti-tumor activity of IL-9R^3x^ pmel T cells was reduced compared to IL-9R^WT^ pmel T cells, with earlier outgrowth of B16-F10 tumor after an initial response. This magnitude of the initial anti-tumor response was also diminished, with complete responses observed in only 1 of 8 mice treated with IL-9R^3x^ pmel T cells compared to 6 of 7 treated with IL-9R^WT^ pmel T cells (Fig. 4G). Together, the detrimental impact of both attenuations and amplifications of IL-9R signaling on anti-tumor efficacy revealed the sensitivity of tumor-specific T cells to variations in the magnitude of signaling and suggests an optimal window in which JAK/STAT signaling can promote anti-tumor activity.

### IL-9R intracellular domain mutants skew STAT phosphorylation profiles and alter in vivo proliferation and anti-tumor efficacy

Given the simultaneous activation of multiple STATs by IL-9R, we explored how the precise stoichiometry of STAT activation, in addition to optimal signal strength, contributes to the superior anti-tumor functions of IL-9R T cells. We confirmed that a Y405F substitution ablated STAT1, STAT3 and STAT4 activation and diminished STAT5 activation (Fig. S5A)^30^. Amino acids immediately adjacent to the phosphotyrosine site can influence STAT-specific binding and activation^31^, so we generated a panel of ten IL-9R mutants, each with a single amino acid substitution of the proline or glutamine residue at the C-terminal side of the tyrosine at position 405 (Y405) (Fig. 5A). T cells engineered with these IL-9R mutants displayed an array of STAT signaling profiles, each with distinct pSTAT stoichiometry (Fig. 5B).

**Figure 5.**
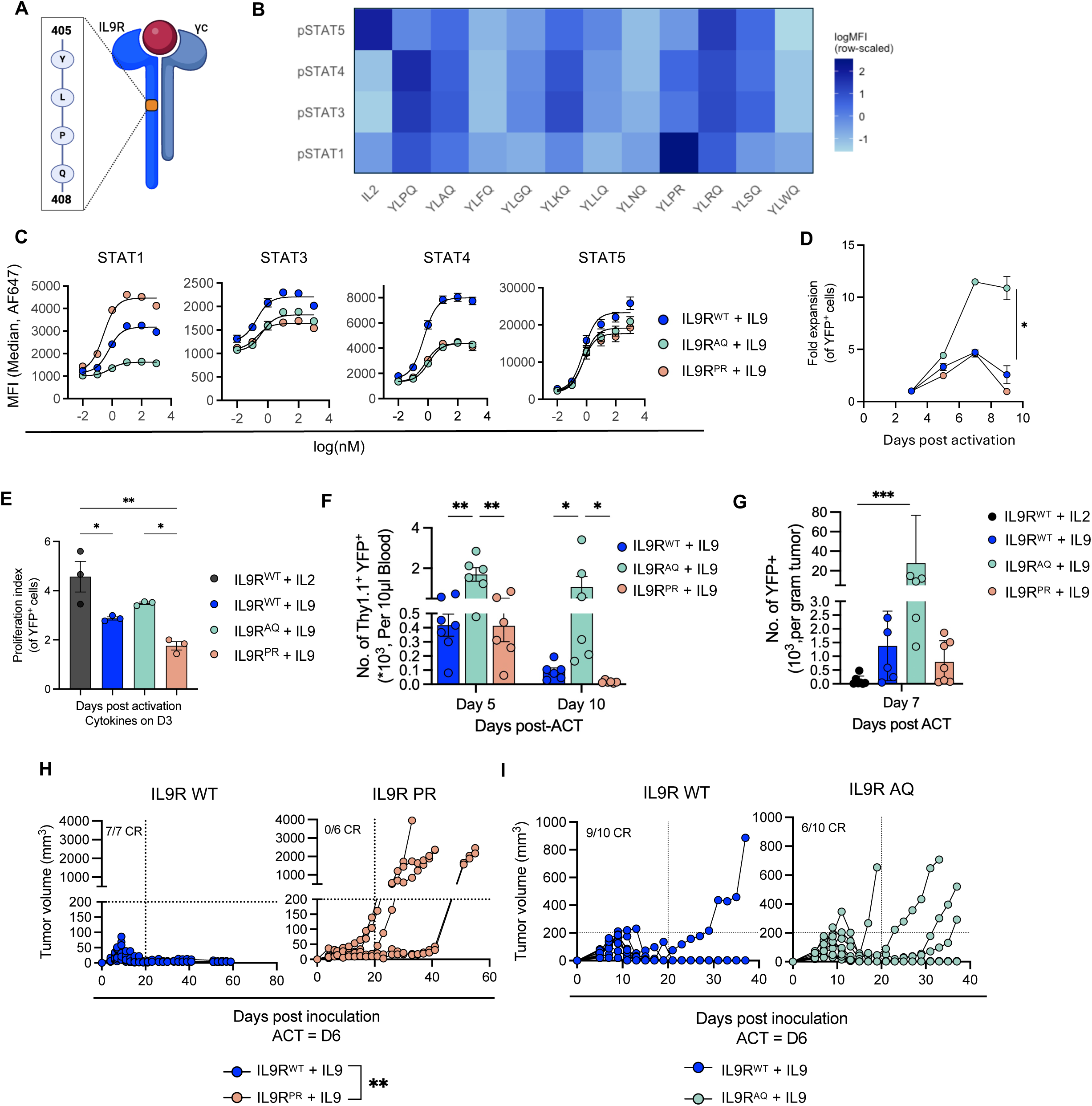
IL-9R intracellular domain mutants skew STAT phosphorylation and alter in vivo proliferative capacity and anti-tumor efficacy. (A) Schematic of the IL-9 receptor complex, highlighting the phosphotyrosine residue within the IL-9R intracellular domain (ICD) and three adjacent amino acids. A panel of ten single amino acid mutations were generated within the ICD at the proline or glutamine residues (created with Biorender.com). (B) Heat map of MFI (log-scaled and row-scaled) at E_max_ for phosphorylation of STAT1, STAT3, STAT4, and STAT5 for C57BL/6 T cells transduced with wildtype IL-9R or one of ten IL-9R mutants. Transduced pmel T cells (technical duplicates) were stimulated with IL-9 for 20 minutes. Data is representative of two independent experiments. (C) Dose-response curves of STAT1, STAT3, STAT4, and STAT5 phosphorylation among transduced IL-9R^WT^, IL-9R^AQ^, and IL-9R^PR^ pmel T cells (YFP+) stimulated with recombinant IL-9 for 20 minutes. Error bars represent SEM of technical duplicates. Data is representative of three biological replicates. (D) Relative in vitro expansion of YFP^+^ IL-9R^WT^, IL-9R^AQ^, IL-9R^PR^ pmel T cells cultured with IL-9 on day 3 post-activation (10 nM; n=3 technical replicates/group). Data is representative of 3 biological replicates. (E) In vitro proliferation index of C57BL/6 T cells engineered with IL-9R^WT^, IL-9R^AQ^, and IL-9R^PR^ over the course of nine days post-activation (transduced on day 1), as measured by dilution of CellTrace Violet dye (n=6 replicates/group). Cells were treated with either recombinant IL-2 or IL-9 (10 nM) on day 3 after T cell activation. Proliferation index was quantified using FlowJo software based on values of histogram peaks (see Figure S5B). (F) Peripheral blood quantification of IL-9R^WT^, IL-9R^AQ^, or IL-9R^PR^ transduced (Thy1.1^+^YFP^+^) pmel T cells in B16-F10 tumor-bearing mice (n=6-7 mice/group) on the indicated days after ACT. IL-9 treatment (5×10^4^ IU i.p., every other day) was started with ACT and continued for 5 doses. Data is representative of at least three independent experiments. (G) Quantification of IL-9R^WT^, IL-9R^AQ^, or IL-9R^PR^ transduced (YFP+) pmel T cells within the tumors (B16-F10) of mice (n=6-7 mice/group) seven days post adoptive cell transfer (ACT). Data is representative of two independent experiments. (H) B16-F10 tumor growth in mice treated with IL-9R^WT^ (n=7 mice) or IL-9R^PR^ (n=6 mice) pmel T cells. Data is representative of at least three independent experiments. (I) B16-F10 tumor growth in mice treated with IL-9R^WT^ or IL-9R^AQ^ pmel T cells (n=10 mice/group). IL-9 treatment as per (F). Data is representative of at least 4 independent experiments. For in vivo experiments, cytokines were tagged with mouse serum albumin (MSA) for half-life extension. *P < 0.05, **P < 0.005, ***P < 0.001, ****P < 0.0001 (two-way ANOVA for D-F; Welch’s t-test for G-I).

We further studied two mutants, YLAQ (IL-9R^AQ^) and YLPR (IL-9R^PR^), that resulted in divergent STAT1 phosphorylation. Compared to IL-9R^WT^, IL-9R^PR^ increased pSTAT1 E_max_ and IL-9R^AQ^ reduced pSTAT1 E_max_ in T cells (Fig. 5C). Notably, IL-9R^AQ^ and IL-9R^PR^ phosphorylated STAT3, STAT4, and STAT5 at similar levels, allowing us to query the impact of T cell intrinsic STAT1 signaling. Compared to IL-9R^WT^, IL-9R^AQ^ and IL-9R^PR^ produced a similar pSTAT5 signal, but weaker pSTAT3 and pSTAT4 signals.

We observed superior in vitro expansion of mouse T cells engineered with IL-9R^AQ^, compared to IL-9R^WT^ (Fig. 5D). There was also a consistent trend toward reduced expansion of IL-9R^PR^ T cells compared to IL-9R^WT^ T cells. The difference in expansion was attributed to an increase in the proliferative index of IL-9R^AQ^ T cells compared to IL-9R^PR^ T cells as measured by dilution of CellTrace Violet dye, with the proliferative index of IL-9R^WT^ T cells falling between IL-9R^AQ^ and IL-9R^PR^ (Fig. S5B-C). We did not observe an increase in apoptotic (Annexin V^+^, Zombie Violet^-^) or dead (Annexin V^+^, Zombie Violet^+^) cells with either mutant (Fig. S5D-E). Given the inverse correlation between STAT1 activity and in vitro proliferation, these results may be attributed to the cytostatic effects of STAT1^32,33^.

The improved proliferation of IL-9R^AQ^ T cells was pronounced in vivo, resulting in ∼5-10-fold greater expansion in the blood between five and ten days after adoptive cell transfer (ACT) compared to IL-9R^WT^ (Fig. 5E). More IL-9R^AQ^ T cells were also observed in the tumor seven days after ACT (Fig. 5F). IL-9R^PR^ T cells expanded less than IL-9R^WT^ in the blood by day 10, a pattern consistent across three independent experiments.

We hypothesized that these differences in proliferative capacity may translate to differences in anti-tumor activity. Compared to treatment of B16-F10 tumor-bearing mice with IL-9R^WT^ pmel T cells (and MSA-IL9), mice treated with IL-9R^PR^ had significantly reduced tumor control, fewer complete responses, and worse survival (Fig. 5G; Fig. S5E), consistent with their diminished proliferative capacity. However, despite the increased peripheral expansion of IL-9R^AQ^ pmel T cells, the tumor growth and survival of mice treated with IL-9R^AQ^ pmel T cells was not superior to mice treated with IL-9R^WT^ pmel T cells (Fig. 5H, Fig. S5F), supporting a decoupling of in vivo expansion and effector function.

### T cell intrinsic STAT1 acts as a rheostat between stem and effector fates of tumor-infiltrating T cells

To further explore the fate of tumor-specific T cells signaling through IL-9R and the biased IL-9R mutants, we performed scRNAseq on tumor-infiltrating pmel T cells engineered with IL-9R^WT^, IL-9R^AQ^ or IL-9R^PR^, eight days after adoptive transfer (Fig. 6A). Consistent with results from Figure 5, tumors in mice treated with IL-9R^WT^ and IL-9R^AQ^ pmel T cells were responding to treatment, whereas tumors in mice treated with IL-9R^PR^ pmel T cells were progressing (Fig. S6A). Likewise, the frequency of engineered (YFP^+^) pmel T cells among infiltrating CD8+ T cells was highest in the IL-9R^AQ^ group (17.0%), followed by IL-9R^WT^ (13.4%) and IL-9R^PR^ (0.7%) (Fig. S6B).

**Figure 6.**
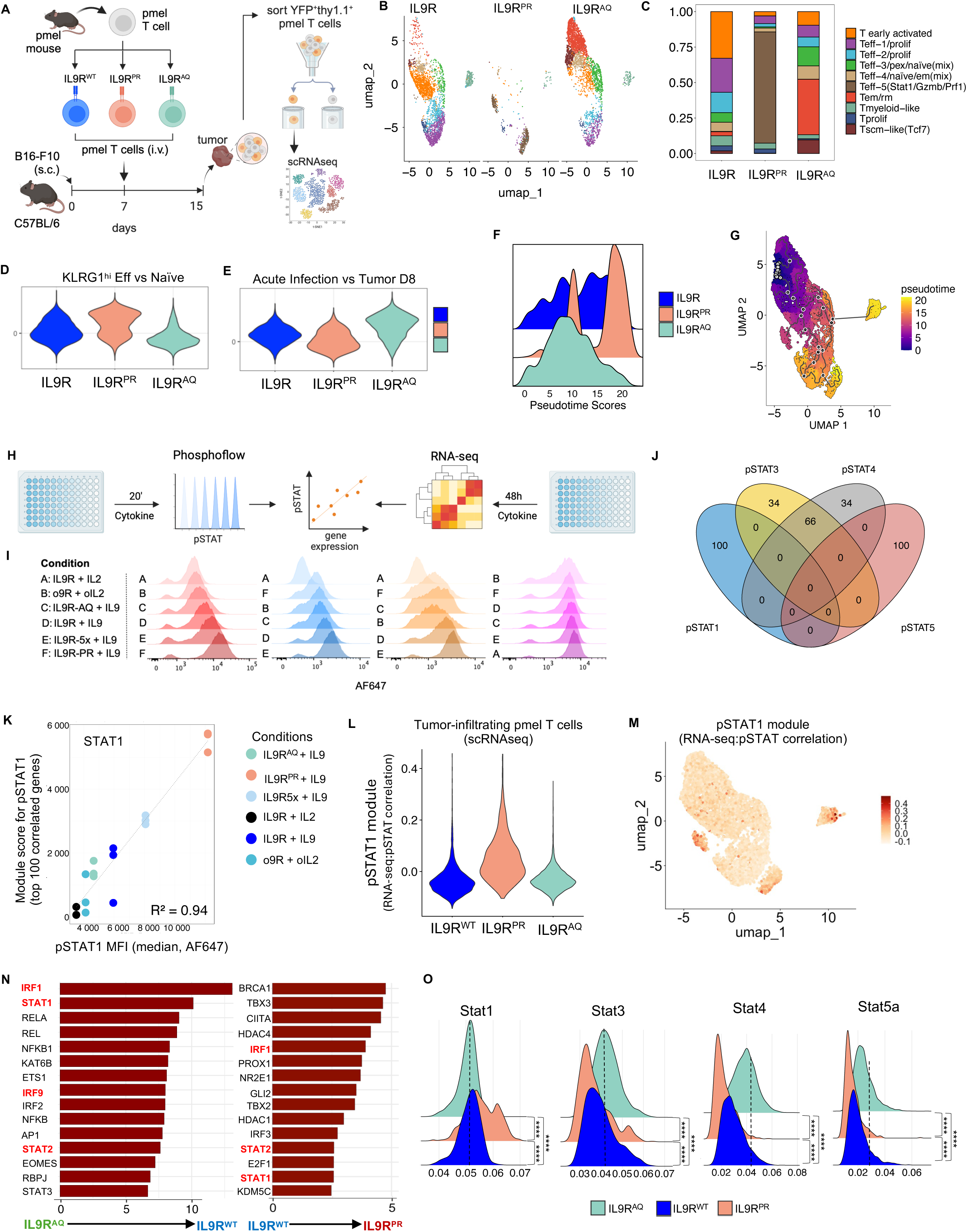
IL-9R variants reveal STAT1 as a rheostat skewing T cells from a stem and memory state toward a terminal effector state. (A) Schematic for in vivo experiment to generate single cell RNAseq data. IL-9R^WT^, IL-9R^AQ^, or IL-9R^PR^ engineered pmel T cells are adoptively transferred into B16-F10 tumor-bearing mice seven days after tumor inoculation. Mice (n=7-8 mice/group) are treated with IL-9 (5×10^4^I.U. every other day) starting on the day of ACT (day 7) until tumors are harvested on day 15. Transduced pmel T cells (Thy1.1^+^YFP^+^) are sorted by FACS prior to library preparation and single cell RNAseq. See related Figure S6A – S6B. (B) UMAP plots based on scRNAseq of n=6,706 cells pmel T cells from (A). Ten major clusters colored according to annotation. See related Figure S6C-E. (C) Propeller plots demonstrating the relative proportion of each cluster from (A), split by treatment group (IL-9R^WT^, IL-9R^AQ^, or IL-9R^PR^). (D) Violin plots summarizing single cell expression of a gene set differentially expressed between KLRG1^hi^ effector and naïve mouse T cells (GSE10239). Gene sets are summarized as a module score in Seurat and plotted by treatment group. (E) Violin plots summarizing single cell expression of a gene set differentially expressed between mouse T cells eight days (D8) after encounter with acute infection versus malignancy (GSE60501). Gene sets are summarized as a module score in Seurat and plotted by treatment group. (F) Ridgeplot of pseudotime scores for single cell data from (A) organized by treatment group. (G) UMAP plot of scRNA-sequencing data from (A) annotated with pseudotime trajectories. Cells within the Tscm-like (Tcf7) cluster were selected as the root for the pseudotime analysis (white circles with black outlines). Black circles with white outlines represent nodes of the differentiation trajectory. (H) Schematic for linking phosphoflow data with RNA-sequencing data. o9R, IL-9R^WT^, IL-9R^AQ^, IL-9R^PR^, or IL-9R^5x^ pmel T cells were treated with cytokines for either 20 mins (phosphoflow) or 48h (RNA-sequencing). MFIs at E_max_ for pSTAT1, pSTAT3, pSTAT4 and pSTAT5 were merged with RNA-sequencing data to identify genes highly correlated with phosphorylation of each STAT protein. (I) Histograms of phosphorylation of STAT1, STAT3, STAT4 and STAT5 for transduced (YFP+) T cells for each condition after 20 minutes of cytokine stimulation at Emax: IL-2 (10nM), IL-9 (10nM) and oIL-2 (10μM). (J) Venn diagram of the top 100 genes most correlated with the phosphorylation of each STAT. (K) Scatterplot depicting the relationship between the pSTAT1 module score (y-axis) and pSTAT1 phosphorylation levels (x-axis) in vitro. Module score was calculated from the expression of 100 genes most strongly correlated with STAT1 phosphorylation in vitro. Shown are biological triplicates for the gene expression data, colored by sample condition. For STAT phosphorylation data, shown are the average of technical replicates. The gray line indicates the linear regression fit. (L) Violin plot depicting projection of pSTAT1 module score from (J)-(K) onto scRNA-seq data from (A), organized by treatment group. (M) Projection of pSTAT1 module score from (J)-(K) onto UMAP of scRNA-seq data from (A). (N) Waterfall plot summarizing transcription factor (TF) enrichment scores when comparing in vitro RNAseq data of IL-9R^WT^ vs IL-9R^AQ^ groups (left) and IL-9R^PR^ vs IL-9R^WT^groups (right). Gene expression changes across IL-9R^AQ^ ➔ IL-9R^WT^➔ IL-9R^PR^ are captured by increasing activity of STAT1 and related TFs (shown in red). (O) Ridgeplots summarizing STAT1, STAT3, STAT4, and STAT5a regulon activity as an AUC score calculated using SCENIC. Results are presented by treatment group, with pairwise statistical comparisons (Wilcoxon Rank-Sum Test).

Among tumor-infiltrating T cells, most cells clustered as subtypes of effector T cells (Fig. S6C, Table S2). These included early activated effectors (T early activated), marked by expression of immune checkpoints *Pdcd1*, *Lag3*, *Havcr2*, and *Vsir*, along with *Gzmb* and *Prf1* expression. Other effector clusters included proliferative effectors (Teff-1/prolif and Teff-2/prolif), and effectors with high expression of *Stat1* (Teff-5). Two clusters consisted of mixed phenotypes of effector, naïve and either precursor exhausted (Teff-3/pex/naive) or effector memory (Teff-4/naïve/em) T cells. Aside from effectors, we identified a larger cluster with features of both effector memory and resident memory T cells (Tem/rm), marked by *Il7r, Itga1, Itga4, Cxcr3, Ccl5* and *Gzma* expression (Fig. S6D). A cluster of stem cell memory-like T cells (T_SCM_-like) had uniquely high expression of *Tcf7,* as well as *Il7r* (Fig. S6D). Finally, we identified clusters of proliferative T cells (Tprolif) and T cells and enriched for a myeloid differentiation gene set (Fig. S6E). Transcription factor and gene regulatory network inference using the Single-Cell rEgulatory Network Inference and Clustering (SCENIC) package identified key regulons for each cluster which aided annotations (Table S2).

Despite the presence of a TCR stimulus within the tumor microenvironment, T cell fate diverged based on the IL-9R state. IL-9R^PR^ pmel T cells, with a high pSTAT1 signal, almost exclusively occupied the Teff-5 (Stat1/Gzmb/Prf1) cluster. IL9R^AQ^ pmel T cells, with a lower pSTAT1 signal, were enriched in Tem/rm and Tscm-like clusters (Fig. 6C, Fig. S6D) and less represented among effector clusters. IL-9R^WT^ pmel T cells occupied an intermediate state, enriched for both early activated effectors (Teff-1/activated) as well as Tem/rm and Tscm-like clusters. Consistent with this gradient of effector phenotype (IL-9R^PR^ > IL-9R^WT^ > IL-9R^AQ^), the overall expression of a gene set distinguishing KLRG1^hi^ effector versus naïve T cells was lowest in IL-9R^AQ^, and highest in IL-9R^PR^ pmel T cells (GSE10239; Fig. 6D)^34^. A similar trend was seen for the expression of *Prf1* and *Gzmb* (Fig. S6F).

We did not find clusters fitting a canonical definition of exhaustion as most cells lacked expression of *Tox* and *Eomes,* consistent with the relatively early timepoint of analysis. Instead, we examined a set of genes that diverge between T cells eight days after exposure to acute infection versus malignancy (GSE60501), a surrogate for a less-differentiated state lacking early features of T cell dysfunction or exhaustion^35^. This gene set was most highly expressed among IL-9R^AQ^ pmel T cells, and lowest among IL-9R^PR^ pmel T cells (Fig. 6E). In summary, compared to IL-9R^WT^, IL-9R^AQ^ drives a memory and stem-like state at the expense of effector differentiation, while IL-9R^PR^ drives a purely effector state with early features of exhaustion.

Pseudotime analysis, anchored by the *Tcf7+* Tscm-like cluster, supported a shared trajectory in which IL-9R^AQ,^ IL-9R^WT^, and IL-9R^PR^ T cells occupy earlier to later stages of differentiation (Fig. 6G). The Tem/rm cluster, comprised primarily of IL-9R^AQ^ pmel T cells, scored low in pseudotime consistent with an earlier differentiation state. In contrast, the Stat1/Gzmb/Prf1 effectors (Teff-5), comprised of IL-9R^PR^ pmel T cells, occupied higher pseudotime scores (Fig. 6G). And while IL-9R^WT^ pmel T cells also consisted of more effectors than IL-9R^AQ^, these were early activated effectors which scored lowest in pseudotime among effector clusters (Fig. 6G).

We hypothesized that the divergent differentiation trajectories were a consequence of pSTAT1, since this was the primary distinguishing signaling feature of IL-9R^AQ^ and IL-9R^PR^ T cells. To explore this, we correlated STAT phosphorylation patterns (measured by phosphoflow) with transcriptomic profiles in vitro (measured by bulk RNA-seq) across a set of six conditions with different patterns of STAT phosphorylation (Fig. 6H-I). For each condition, T cells were engineered with a cytokine receptor (o9R, IL-9R, IL-9R^AQ^, IL-9R^PR^ or IL-9R^5x^) and treated with cytokine (IL-2, IL-9, or oIL-2) for 20’ to evaluate signaling by phosphoflow and for 48h to evaluate their transcriptomic profile by RNA-seq. We identified the top 100 genes correlated with phosphorylation of each STAT protein, which for pSTAT1 and pSTAT5 were entirely unique (Fig. 6J, Table S3). Sixty-six of 100 genes most correlated with pSTAT3 and pSTAT4 were shared. The highest correlation between gene expression and STAT phosphorylation was observed for pSTAT1, reflecting the range and even distribution of pSTAT1 activation across the sample set (Fig. 6K).

We generated a gene expression module based on the genes most correlated with phosphorylation of each STAT, which were then projected onto the scRNA-seq data from Fig. 6A. The pSTAT1 module was the dominant feature distinguishing IL-9R^PR^ pmel-1 T cells from IL-9R^WT^ and IL-9R^AQ^ pmel-1 T cells, consistent with the high pSTAT1 levels upon IL-9R^PR^ activation and validating the fidelity between the in vitro and in vivo transcriptomic effects of IL-9R signaling (Fig. 6K-L). We did not find differences in pSTAT3, pSTAT4, or pSTAT5 gene modules between the groups, though these are less sensitive for detecting differences given the lower correlation between gene expression and STAT phosphorylation.

Since the high pSTAT1 signal of IL-9R^PR^ T cells was driving the terminal effector differentiation in vivo, we wished to understand if the low pSTAT1 signal of IL-9R^AQ^ T cells contributed to their less-differentiated stem-like and memory state compared to their IL-9R^WT^ counterparts. As noted above, in addition to higher STAT1 phosphorylation, IL9R^WT^ signaling also induces higher STAT3 and STAT4 phosphorylation compared to IL-9R^AQ^. Based on in vitro RNA-seq, we inferred differential transcription factor (TF) activity and identified STAT1 and related TFs (IRF1, IRF2 and IRF9), as the dominant distinguishing feature between IL-9R^WT^ and IL-9R^AQ^ T cells (Fig. 6N). SCENIC analysis of the scRNA-seq dataset also demonstrated a modest increase in the STAT1 regulon in tumor-infiltrating IL-9R^WT^ compared to IL-9R^AQ^ pmel T cells (Fig. 6O). In fact, among STAT1, STAT3, STAT4 and STAT5, STAT1 was the only regulon enriched in IL-9R^WT^ compared to IL-9R^AQ^ pmel T cells. Furthermore, genes differentially expressed by IL-9R^AQ^ T cells were enriched for genes from Hallmark IL2/STAT5 signaling and IL6/JAK/STAT3 signaling gene sets (adjusted p-value = 0.001 and 0.07, respectively; Fig. S6J).

We also noted that STAT3, STAT4 and STAT5 regulons were enriched in tumor-infiltrating IL-9R^AQ^ pmel T cells compared to both IL-9R^WT^ and IL-9R^PR^ pmel T cells. The enrichment of STAT3 and STAT5 regulons is consistent with the stem, memory^28,36^ and proliferative qualities^27,37^ of IL9R^AQ^ pmel T cells (Fig. 6N). However, this is also partly counterintuitive considering that IL-9R^AQ^ results in stronger pSTAT3 and pSTAT4 activation (and similar pSTAT5 activation) compared to IL-9R^WT^. We speculate that differences between STAT profiles based on proximal signaling versus in vivo transcriptomics may relate to complex downstream interplay and competition between STAT proteins (see discussion).

In summary, we find divergent fates of tumor-infiltrating antigen-specific T cells with activated signaling through IL-9R^WT^, IL-9R^AQ^, or IL-9R^PR^. T cell intrinsic STAT1 activity underlies the differentiation trajectory from a less-differentiated, stem-like state (IL-9R^AQ^, STAT1-low) to a terminally differentiated effector (IL-9R^PR^, STAT1-hi), with IL-9R^WT^ (STAT1-mid) occupying a middle ground encompassing both stem-like and effector states.

### Potency and sensitivity of proximal IL-9R signaling for human CAR T cells

Similar to results in the mouse system, human T cells engineered with IL-9R and cultured with IL-9 phosphorylate STAT1, STAT3, and STAT5 at higher levels than human T cells engineered with the human orthogonal chimeric o9R (ho9R) and cultured with the human orthogonal IL-2 (hoIL-2) (Fig. 7A), and also phosphorylate STAT4. We then co-engineered human T cells with a second-generation chimeric antigen receptor (CAR) targeting CD19 (CD19-BBz) and the human IL-9R (hIL-9R; Fig. S7A). In a repetitive killing assay with NALM6 tumor cells which mimics chronic antigen stimulation, CD19-BBz CAR T cells co-transduced with IL-9R and cultured with IL-9 outperformed the same cells cultured with no cytokine or IL-2 between five and nine rounds of tumor killing (Fig. S7B). We then treated Nalm6 leukemia-bearing NSG mice with CD19-BBz CAR T cells co-engineered with or without hIL-9R, as well as human IL-9 or IL-2 (Fig. 7B). Mice treated with IL-2 were euthanized due to rapid weight loss indicating toxicity of IL-2. In contrast, IL-9 did not cause weight loss (Fig. 7C). While CD19-BBz CAR T cells had only a modest anti-tumor effect, dual expressing CD19-BBz hIL-9R T cells (in mice treated with IL-9) resulted in prolonged tumor control and survival (Fig. 7D-E). IL-9 did not produce anti-tumor effects in the absence of IL-9R-engineered T cells.

**Figure 7.**
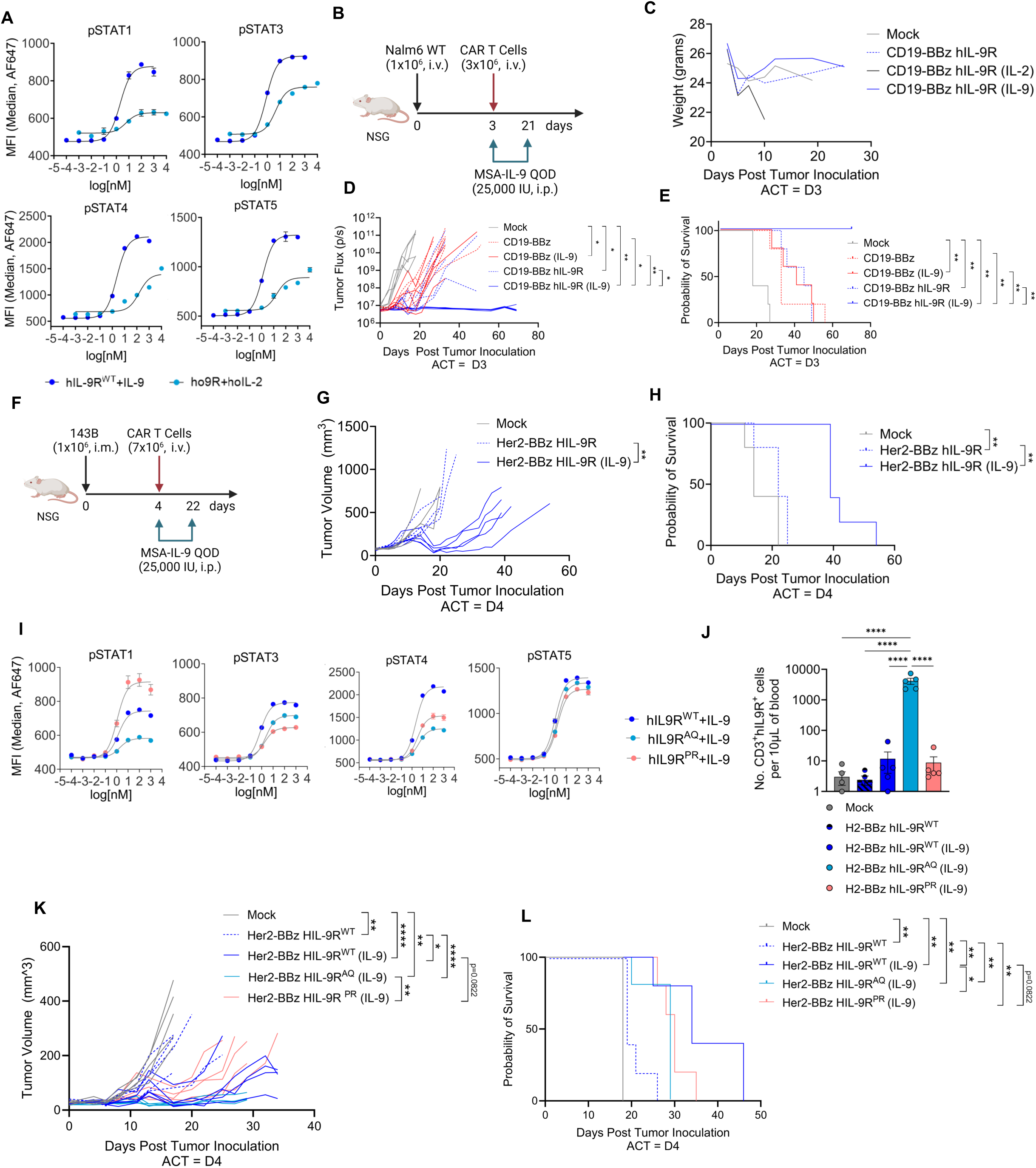
The balance of JAK/STAT signaling through IL-9R and IL-9R variants, and their impact on anti-tumor activity, is conserved in human CAR T cells. (A) Dose-response curves of STAT1, STAT3, STAT4, and STAT5 phosphorylation in human T cells transduced with human IL-9R or human o9R (YFP+) and stimulated with either human IL-9 or human oIL-2 for 20 minutes. Data are representative of two independent donors. (B) Schematic of Nalm6 human leukemia orthotopic model in NSG mice treated with second generation human CAR T cells targeting CD19 (CD19-BBz) and co-transduced with human IL-9R. Mice were also treated with either human IL-2 or IL-9 (or no cytokine). (C) Weight of NSG mice (n=5 mice/group) in response to treatment schema shown in (B). Mice were weighed twice per week. Mice were euthanized at day 10 in the group receiving IL-2 due to toxicity. (D) Bioluminescence (photons/second) of tumors in mice treated as described in (B). (E) Survival curves of mice from (D). (F) Schematic of 143B human osteosarcoma orthotopic solid tumor model in NSG mice treated with second generation human CAR T cells targeting Her2 (Her2-BBz). CAR T cells were cotransduced with human IL-9R (or not) and treated with human IL-9 (or not). (G) Tumor growth as measured by leg volume of mice (n=5 mice/group) as treated per (F). (H) Survival curves of mice from (G). (I) Dose-response curves of STAT1, STAT3, STAT4, and STAT5 phosphorylation in human T cells transduced with human IL-9R, IL-9R^PR^ or IL-9R^AQ^ and stimulated with IL-9 for 20 minutes. Shown are technical duplicates (bars represent SEM) and representative of two independent donors. (J) In vivo expansion and enrichment of CAR T cells from (K). Panel shows absolute number of (YFP^+^) CAR T cells per 10 microliters of blood twelve days after tumor inoculation (8 days after ACT). (K) Tumor growth as measured by leg volume of mice (n=5 mice/group) treated as per schematic in (F). Tumor volume was measured three times per week. (L) In vivo survival curves of mice from (K). For all experiments in Figure 7, cytokines were tagged with mouse serum albumin for half-life extension.*P < 0.05, **P < 0.005, ***P < 0.0005, ****P < 0.0001 (two-way ANOVA for D, G, J; Mantel-Cox test for E, H, K; one-way ANOVA for L).

We then evaluated this approach against 143B, a solid tumor human osteosarcoma model inoculated orthotopically into the hind leg of NSG mice (Fig. 7F). Second generation CAR T cells targeting Her2 (H2-BBz) had minimal anti-tumor effect alone (Fig. 7G-H). However, dual expressing H2-BBz hIL9R IL-9RT cells in mice treated with IL-9 resulted in superior anti-tumor activity and prolonged survival (Fig. 7G-H).

We also found that the signaling patterns of mutant IL-9 receptors, IL-9R^AQ^ and IL-9R^PR^, were conserved between mouse and human systems (Fig. S7C-D). As in the mouse system, IL-9R^AQ^ reduced STAT1 phosphorylation, whereas IL-9R^PR^ increased STAT1 phosphorylation (Fig. 7I). Human IL-9R^AQ^ and IL-9R^PR^ produced comparable pSTAT3 and pSTAT4 signals, and both mutants reduced STAT3 and STAT4 phosphorylation relative to IL-9R^WT^. Additionally, pSTAT5 levels were similar between IL9R and the mutants.

Likewise, the biological effects of the mutant IL-9 receptors were conserved between mouse and human systems. Compared to H2-BBz CAR T cells co-engineered with IL-9R^WT^, cells co-engineered with IL-9R^AQ^ expanded more in vitro, whereas IL-9R^PR^ cells expanded less. In vivo, the expansion and enrichment of CAR T cells co-engineered with IL-9R^AQ^ was superior to CAR T cells co-engineered with IL-9R^WT^ or IL-9R^PR^ (Fig. 7J, Fig. S7E). We did not observe a significant difference between in vivo expansion of IL-9R^WT^ and IL-9R^PR^ CAR T cells, suggesting that CAR signaling may counteract the anti-proliferative effects of IL-9R^PR^ signaling. Treatment with IL-9 significantly improved tumor control in mice treated with CAR T cells co-engineered with IL-9R^WT^ and IL-9R^AQ^, but not IL-9R^PR^ (Fig. 7K), consistent with the inferior anti-tumor functions of mouse IL-9R^PR^ pmel T cells. Treatment with IL-9R^WT^ CAR T cells prolonged survival compared to IL-9R^PR^ (p=0.078) and IL-9R^AQ^ CAR T cells, though survival in mice treated with IL-9R^AQ^ CAR T cells was confounded by deaths related to toxicity presumably related to rigorous CAR T cell expansion. Altogether, the pattern of IL-9R signaling, the signaling effects of the single amino acid mutations, and the resulting in vivo expansion and anti-tumor efficacy mirror the results in the mouse system, further supporting its role as a driver of anti-tumor functions in T cells that is exquisitely sensitive to its optimal balance of JAK/STAT signaling.

## DISCUSSION

Despite significant advancements, including two FDA approvals^38,39^, the widespread adoption of T cell therapies for solid tumors faces substantial challenges. Treatments require large-scale adoptive transfers and toxic lymphodepletion regimens and, even then, T cells are limited by poor trafficking, chronic antigen exposure and hostile tumor microenvironments. While overexpressing or knocking out genes in T cells are potential solutions^40,41^, these lack on/off switches and raise safety concerns. Cytokines present an attractive alternative by providing a regulatable system to modulate engineered T cells in vivo through coordinated signaling pathways. IL-2 is used with tumor-infiltrating lymphocyte (TIL) therapy, but severe toxicities limit its therapeutic potential^3^. Similar concerns remain for other γ_c_ cytokines, as well as IL-12 and IL-18^42–44^.

Here, we report that IL-9 stands out among γ_c_ cytokines because its private receptor (IL-9R) has remarkably restricted expression across tissues (Fig. 1). This explains how high doses of half-life extended IL-9 are well-tolerated by mice (Fig. 2A-E) and systemic IL-9 administration induces negligible gene expression changes in lymph nodes (Fig. 1E). Consistent with this finding, a companion article published in this issue (Castelli, et al.) reported that tumor-directed IL-9 signaling only altered IL-9R-engineered CAR T cells without impacting the composition of the tumor microenvironment.

This is not to suggest that IL-9 does not have important roles driven by rare IL-9R expressing cells in specific biological contexts^45^. IL-9 has been associated with type 2 immune responses, particularly in allergic conditions or asthma^46–48^ and also contributes to protective immunity against parasitic helminths^49^. The IL-9/IL-9R axis contributes to various immunological processes, including mast cell activation, ILC2 function and B and T cell differentiation. In airway inflammation and hyperresponsiveness, IL-9 can act as a biological response amplifier^48,50,51^. Even so, it is notable that IL-9 deficient mice are healthy with a normal immunophenotype, including normal T cell differentiation and activation, normal antibody responses, normal responses to parasitic infection, and normal Th2 response to allergens in the lung^52,53^. And based on population-level whole exome sequencing data, IL-9 scored lowest among a panel of canonical cytokines with respect to indispensability to evolutionary fitness (Fig. 1F).

We took advantage of the paucity of IL-9R expression to engineer T cells with the native IL-9R. Coadministration of IL-9 with as few as 0.5×10^6^ tumor-specific T cells engineered with IL-9R generated robust anti-tumor responses in mice without preconditioning lymphodepletion, outperforming our benchmark o9R system (Fig. 2F-G). These findings were consistent across multiple mouse and human models, as well as data from Castelli, et al (REF). IL-9R T cells demonstrate superior signaling, transcriptomic, and biological outputs compared to the synthetic o9R system. This is consistent with the low nanomolar affinity of the IL-9/IL-9R interaction, a distinct advantage over synthetically designed orthogonal systems (e.g. o9R), which are inherently engineered to be low affinity systems with weaker signal strength^8,11^.

The stronger signal through IL-9R revealed the unexpected phosphorylation of STAT4 by IL-9 (Fig. 3C-D). The canonical activator of STAT4 in T cells, IL-12, is well known for promoting anti-tumor T cell functions^23^, but its use has been limited by toxicity risks^43^. IL-9 creates an avenue to deliver potent STAT4 signals to engineered T cells without toxicity, while also retaining STAT5, which promotes T cell proliferation and functional effector states^27,54^, and STAT3, which can promote stemness^28^ (Fig. 3A).

Our findings reveal a “goldilocks phenomenon” with respect to the JAK/STAT signaling strength of IL-9R. Either attenuation through a mutation at the γ_c_ interface of IL-9 (Q115T) (Fig. 4B-D) or amplification via repeated phosphotyrosine ICD motifs (IL-9R^3x^ and IL-9R^5x^) (Fig. 4E-F) proved detrimental to anti-tumor efficacy. Likewise, T cells were exquisitely sensitive to changes in pSTAT stoichiometry generated through single amino acid substitutions adjacent to the primary phosphotyrosine site of the IL-9R ICD^55^. We explored two IL-9R mutants with divergent pSTAT1 signals (IL-9R^AQ^ and IL-9R^PR^) to clarify the role of T cell intrinsic STAT1 in anti-tumor functions (Fig. 5). STAT1 has anti-proliferative effects in T cells^32,33^, but has also been linked to clonal expansion, memory formation^56^, and effector function. In our model, amplification of pSTAT1 (IL-9R^PR^) diminished expansion and reduced anti-tumor efficacy, confirming its anti-proliferative role (Fig. 5D-G). Attenuation of pSTAT1 (IL-9R^AQ^) resulted in superior expansion, particularly in vivo, accompanied with better tumor infiltration. However, superior expansion did not translate to better anti-tumor efficacy (Fig. 5H), demonstrating that T cell expansion in vivo can be uncoupled from anti-tumor efficacy.

Subtle shifts in pSTAT stoichiometry have potent effects on cell fate, even in the presence of an active TCR stimulus. Transcriptomic differences driven by IL9R variants were captured along a pSTAT1 axis, with high pSTAT1 driving the development of a terminally differentiated effector state (IL-9R^PR^), low pSTAT1 yielding a proliferative, stem-like state (IL-9R^AQ^), and IL-9R^WT^ occupying an intermediate state. This is consistent with previous work demonstrating that interferon signaling, through IRF2, promotes CD8+ T cell exhaustion^57^, and also provides a plausible explanation for why JAK/STAT inhibition in the context of chronic interferon signaling may restore T cell effector functions^58^. However, pSTAT1 may not be entirely detrimental to anti-tumor immunity: lower pSTAT1 signals (IL-9R^AQ^) restrained effector differentiation, and IL-9R^AQ^ pmel T cells did not improve anti-tumor efficacy despite superior T cell expansion.

The reduced pSTAT1 signal produced by IL-9R^AQ^ was also associated with an enrichment in STAT3 and STAT5 activity inferred from single cell transcriptomic data, consistent with their stem, memory^28,36^ and proliferative qualities^27,37^ of IL9R^AQ^ pmel T cells (Fig. 6N). However, this was partly unexpected because IL-9R^AQ^ produces a weaker pSTAT3 signal than IL-9R^WT^ by phosphoflow (and similar pSTAT5 signal). We speculate that the discrepancy between STAT activity assessed by phosphorylation and STAT activity inferred from scRNA-seq data may be related to downstream competition between simultaneously active STAT proteins, a phenomenon described in the fate of Th17 cells^59^. The weaker pSTAT1 signal of IL-9R^AQ^ may result in less competition with pSTAT3, such that even a relatively weak pSTAT3 signal generates a stronger downstream biologic effect.

In summary, our study uncovers and leverages the uniquely restricted normal tissue expression of IL-9R. The IL-9/IL-9R is a natural cytokine-receptor pair with orthogonal qualities, and in models of engineered T cell therapy, outperforms its synthetic counterpart (o9R). The activity of IL-9R can be linked to its higher affinity and more potent signaling, which unexpectedly includes STAT4, not previously considered a primary target of γ_c_ signaling. The highly potent anti-tumor functions of T cells signaling through IL-9R – validated in multiple mouse and human CAR T cell models against hematologic and solid tumors – can be traced to an optimal strength and balance of IL-9R signals, with pSTAT1 tuning cells between a stem-like and terminal effector state.

### Limitations of the Study

Our evidence supports the hypothesis that systemic IL-9 administration would be safe in patients; ultimately, however, the safety of IL-9 in humans will be determined in phase 1 clinical trials. With respect to the effects of manipulating JAK/STAT signaling, our results must be considered in the context of the JAK/STAT profile of the IL-9R and may not be generalizable to other contexts where STAT1, STAT3, STAT4, and STAT5 are not simultaneously active. Future studies will be necessary to interrogate downstream mechanisms of signaling competition and cooperation between simultaneously phosphorylated STATs. Lastly, the in vivo studies of human T cells in NSG mice may not fully capture the effects of JAK/STAT pathway manipulations, which can have discrete effects on interactions with host immune cells.

## METHODS

### Protein production

DNA encoding mouse and human IL-2, mouse and human IL-9, and mouse IL-9^Q115T^ were cloned into the mammalian expression vector pD649, which includes a C-terminal 8xHis tag for affinity purification. DNA encoding mouse serum albumin (MSA) was purchased from Integrated DNA Technologies (IDT) and cloned into pD649 as an N-terminal fusion. Mammalian expression DNA constructs were transfected into Expi293F cells using the Expi293 Expression System (Thermo Fisher Scientific) for secretion and purified for the clarified supernatant by nickel affinity resin (Ni-IMAC, Thermo Fisher Scientific) followed by size-exclusion chromatography with a Superdex-200 column (Cytiva) and formulated in sterile phosphate-buffered saline (PBS) for injection. Endotoxin was removed using the Proteus NoEndo HC Spin column kit following the manufacture’s recommendations (VivaProducts) and endotoxin removal was confirmed using the Pierce LAL Chromogenic Endotoxin Quantification Kit (Thermo Fisher Scientific). Proteins were concentrated and stored at −80°C until use.

### Mammalian Expression Vectors

cDNA encoding mouse orthogonal IL-2Rb and geneblock cDNA encoding mouse ICDs of IL-9R (IDT) were cloned into the retroviral vector pMSCV-MCS-IRES-YFP by PCR and isothermal assembly (ITA). Human orthogonal IL2Rb-ECD-IL9R-ICD (ho9R) were similarly cloned into the pMSCV vector. cDNA encoding the native mouse and human IL9R-P2A-YFP were synthesized (VectorBuilder) and cloned into the pMSCV vector. CD19-4-1BBζ and HER2-4-1BBζ constructs were generous gifts from Crystal Mackall.^60^

Single amino acid mutations of the IL-9R, including those at and near Y405 (mouse) and Y407 (human), and the single amino acid mutation of IL-9 (Q115T), were made through Q5 site-directed mutagenesis (New England Biolabs). DpnI-treated plasmids were subsequently transformed using DH5a competent cells (New England Biolabs) and grown on LB agar plates containing 100 μg ml^-1^ ampicillin overnight. Bacterial colonies were extracted (Zymo Research) and plasmids were subsequently screened and sent for full plasmid sequencing (Azenta) for validation. To generate IL-9R receptors with repetitive phosphotyrosine motifs (IL-9R^3x^ and IL-9R^5x^), geneblocks encoding amino acids from position 386 to position 425 were repeated three times or five times in tandem, followed by the remainder of the IL-9R intracellular domain.

### Mice

Mice were housed in animal facilities approved by the Association for the Assessment and Accreditation of Laboratory Care. Five- to six-week-old C57BL/6 (C57BL/6J) mice were purchased from Jackson laboratory. Five- to eight-week-old pmel-1 TCR/Thy1.1 (pmel; B6.Cg-Thy1a/Cy Tg(TcraTcrb)8Rest/J) transgenic mice were originally purchased from the Jackson laboratory and maintained in the Stanford University-Research Animal Facility (RAF) Facility. NOD.Cg-Prkdcscid Il2rgtm1Wjl/SzJ (NSG) mice were originally purchased from the Jackson Laboratory and maintained in the Stanford University Lorry Lokey (SIM1) facility by the Stanford Veterinary Service Center before transfer to the Stanford University Comparative Medicine Pavilion for experimental procedures.

### Cell lines and cell culture

The B16-F10 mouse melanoma cell line was purchased from ATCC and cultured with RPMI 1640 with GlutaMAX (Thermo Fisher Scientific) containing 10% fetal bovine serum (FBS, Omega Scientific) and penicillin-streptomycin (100 µg ml^-1^, Thermo Fisher Scientific) (RPMI-C). The KP-gp100 mouse soft-tissue sarcoma cell line was derived from KP3172, a tumor cell line derived from a Cre-induced sarcomas arising in the hind leg of KP (Kras^LSL-G12D^, p53^flox^) mice. KP317 was retrovirally transduced with a construct containing mouse gp100 with mutations KVP at amino acids 25-27 to generate KP-gp100; the cell line was maintained in RPMI-C. Primary mouse T cells derived from C57BL/6 or pmel transgenic mice were cultured in RPMI-1640 with GlutaMAX supplemented with 10% FBS, penicillin-streptomycin, 50 μM 2-mercaptoethanol (Gibco), 1x non-essential amino acids (Gibco), 1x sodium pyruvate (Gibco), 20 mM HEPES (Gibco), and 1x GlutaMAX. HEK293T cells were purchased from ATCC and cultured in high-glucose DMEM (Gibco) supplemented with 10% FBS, 1x GlutaMAX, 1x sodium pyruvate), and penicillin-streptomycin (DMEM-c). HEK293GP cells were purchased from ATCC and cultured in high-glucose DMEM (Gibco) supplemented with 10% FBS, 2mM GlutaMAX, 10mM HEPES, and penicillin-streptomycin (DMEM-c). Expi293F cells were purchased from Thermo Fisher Scientific and cultured in Expi293 Expression Medium (Thermo Fisher Scientific). Nalm6-FFluc cells were a generous gift from Crystal Mackall and cultured in RPMI (Gibco) supplemented with 10% FBS, 2mM GlutaMAX, 10mM HEPES, and penicillin-streptomycin (RPMI-c). 143B-GFP-FFluc cells were a generous gift from Crystal Mackall and cultured in DMEM-c. Cell lines were periodically authenticated (ATCC) and periodically tested for mycoplasma infection using a mycoplasma detection kit (Vazyme).

### Retrovirus production

For production of retrovirus to engineer mouse T cells, HEK293T (ATCC) cells were seeded at 4.5 x 10^5^ cells per well in a 6-well tissue culture treated plate. 6-well tissue culture treated plates were coated with 0.01% poly-l-lysine (Sigma-Aldrich) prior to seeding. Following an overnight incubation, the medium was replaced with pre-warmed DMEM-c. For each well of transfection, 2.5 µg of plasmid (1.25 µg of pMSCV retroviral vector plus 1.25 µg of pCL-Eco packaging vector) was added to 250 µl of Opti-MEM I Reduced Serum Medium (Thermo Fisher Scientific), followed by 5.5 µl of Lipofectamine™ 3000 reagent (Thermo Fisher Scientific) and 5 µl of P3000™ enhancer reagent, with gentle vortexing. After a 15-minute incubation at 25°C, the transfection mixture was gently added to HEK293T cells and cultured overnight. After 18 to 20 h, the medium was replaced with fresh DMEM-c with 20 mM HEPES and cultured for another 24 h before collection at 48 hours post-transfection. The medium was collected and filtered through a 0.22 µM Whatman filter (Cytiva). Transfection was visually confirmed through confocal microscopy (Leica) for YFP marker. If not used immediately, the virus was frozen for storage in −80°C.

For production of retrovirus to engineer human T cells, HEK293GP (ATCC) cells were cultured in complete DMEM (DMEM-c) media (10% FBS, 100 U/mL penicillin, 100 μg/mL streptomycin, 2mM GlutaMAX, 10mM HEPES). Prior to plating of HEK293GP cells, 10cm tissue-culture treated dishes were coated with 5 mL of 0.01% poly-L-lysine for at least 10 minutes. Cells were subsequently plated at a density of 6.5 x 10^6^ ml^-1^ per dish in 10 mL total DMEM-c and incubated overnight at 37°C. After 24 hours, transfection was performed using 9 μg vector plasmid, 1.5 mL of Mix A (1.5 mL Opti-Mem, 4.5 μg RD114), and 1.5 mL of Mix B (1.5 mL Opti-MEM, 33uL Lipofectamine 3000, and 2uL P3000 per μg vector plasmid) per plate. 24 hours after transfection, fresh media was replenished. Virus was collected at 48- and 72-hours post-transfection. If not used immediately, the virus was frozen for storage in −80°C.

### Activation and retroviral transduction of primary mouse T cells

For retroviral transduction of mouse T cells, splenocytes from the five- to ten-week-old mice were mechanically digested and filtered through a 70 µM strainer (Fisher Scientific). Red blood cells were lysed with eBioscience RBC Lysis Buffer (Thermo Fisher Scientific) for 5 minutes at 4°C. Splenocytes were resuspended in PBS with 2% FBS and 1 mM Ethylenediaminetetraacetic acid (EDTA, Thermo Fisher Scientific), and enriched for CD3^+^ cells by magnetic bead separation, according to the manufacturer’s protocol (StemCell). C57BL/6-derived mouse T cells were activated with a 1:1 bead:cell ratio of mouse CD3/28 Dynabeads™ (Thermo Fisher Scientific) in fresh T cell medium with 100 U ml^-1^ rmIL-2 overnight. On day of transduction, CD3/28 Dynabeads™ were removed prior to transduction by magnetic bead separation. Isolated pmel T cells were activated with 100 U ml^-1^ recombinant mouse IL-2 (rmIL-2) (Peprotech) and 1 µg ml^-1^ human gp100 peptide (Anaspec) the day before transduction. One day before transduction, six-well non-tissue culture treated plates (Thermo Fisher Scientific) were coated with 37.5 µg ml^-1^ retronectin (Takara Bio) in PBS per well and placed at 4°C overnight. The following day, retronectin-coated plates were blocked with 0.5% BSA (Sigma-Aldrich) in PBS for 30 minutes at 25°C and washed out with PBS. Viral supernatant (2 ml) was added to each well and spun at 2000*g* for two hours at 32°C without brake. Viral supernatant was carefully removed, 3 ml of activated T cells (1 x 10^6^ ml^-1^) were added to each well with 100 U ml^-1^ rmIL-2, and spun at 2000*g* for 10 minutes at 32°C without brake. Cells were then cultured for 18-22 h at 37°C. Cells were collected via gentle pipetting and resuspended at 1 x 10^6^ ml^-1^ in fresh mouse T cell medium with 100 U ml^-1^ rmIL-2 and expanded overnight before further downstream cellular assays. For in vivo tumor assays, transduced cells were used either immediately after collection or one day post-transduction. YFP served as a surrogate marker for the expression of all mouse constructs used and was checked on day of use.

### Isolation of PBMC and T cells

Healthy donor Leukocyte Reduction System (LRS) chambers were purchased from the Stanford Blood Center according to an institutional review board (IRB)-exempt protocol. Peripheral blood mononuclear cells (PBMCs) were isolated using Ficoll-Paque Plus (Cytiva) density gradient centrifugation and frozen in a Mr. Frosty™ Freezing Container in aliquots of 10 x 10^6^ PBMCs mL^-1^ of CELLBANKER 2 serum-free cell cryopreservation medium (AMSBIO). Aliquots were stored frozen at −80°C

### Activation and retroviral transduction of primary human T cells

Human T cells were isolated from cryopreserved PBMCs using the EasySep™ Human T Cell Isolation Kit (StemCell) according to the manufacturer’s protocol. Human T cells were activated using Dynabeads™ Human T-Activator CD3/CD28 (Thermo Fisher Scientific) for T Cell expansion and activation at a 3:1 ratio of beads:cells and cells were incubated at 37°C for three days at a concentration of 1 x 10^6^ cells ml^-1^. Cells were cultured in complete AIM-V media (10% fetal bovine serum, 100 U/mL penicillin, 100 μg/mL streptomycin, 2mM GlutaMAX, 10mM HEPES (Thermo Fisher Scientific) and 100 U/mL recombinant human IL-2; Peprotech). Transduction was performed on days 3 and 4 after activation. Twelve-well, non-tissue culture treated plates (Thermo Fisher Scientific) were coated with 18.75 µg mL^-1^ retronectin (Takara Bio) in PBS per well and left at 4°C overnight or for two hours at 25°C. On the day of transduction, the retronectin was transferred to a new 12-well non-tissue culture treated plate, which was subsequently stored at 4°C overnight. The original retronectin-treated plate was then blocked with 1 mL of 1% bovine serum albumin (BSA; Sigma-Aldrich) in PBS per well for 10-20 minutes. Afterward the BSA was removed and 1 mL of 48 hr retroviral supernatant was added per well. For co-transductions, 1 mL of supernatant per construct was added per well (for a total of 2 mL per well). Following the addition of viral supernatant, the plate was spun at 2000 x g for 2 hours at 32°C. Subsequently, viral supernatant was aspirated, and the activated Day 3 T cells were added at a concentration of 0.5 x 10^6^ cells ml^-1^ in 1 mL per well. The plate was then incubated at 37°C overnight. The transduction was repeated the following day using 72 hr supernatant. On Day 5 post activation, the DynaBeads were magnetically removed from T cells and cells were resuspended in fresh media at a concentration of 0.5 x 10^6^ cells ml^-1^. On Day 7 post activation, T cells were passaged and resuspended in fresh media at a concentration of 0.5 x 10^6^ cells ml^-1^. On Day 10, transduction efficiency was assessed by flow cytometry and T cells were used in subsequent assays.

### In vivo mouse tumor studies

#### Syngeneic mouse tumor studies

For in vivo tumor growth experiments, early-passage cell lines were used (fewer than 10 passages). B16-F10 or KP-gp100 tumor cells (5 x 10^5^ cells) were resuspended in 100 µl of PBS and injected subcutaneously in the right flank of 6–10-week-old C57BL/6J mice. Prior to adoptive cell transfer, mice were randomized based on average tumor size. 2 x 10^6^ – 5 x 10^6^ non-sorted, transduced T cells (derived from pmel mice) were adoptively transferred five or six days after tumor inoculation unless otherwise indicated. Specifically, T cells were resuspended in 50 µl of PBS per mouse and administered by retroorbital intravenous injection. Where indicated, mice received treatment with cytokines: mouse serum albumin (MSA)-bound mouse IL-2 (MSA-IL2), MSA-IL9, MSA-IL9^Q115T^, MSA-oIL2 (5 x 10^4^ units per dose, intraperitoneal) for five doses (or longer, where indicated), administered every other day starting on the day of adoptive cell transfer (unless otherwise indicated). Tumor size (length x width) was measured with calipers three times a week and volume was calculated as (length x width^2^)/2). Peripheral blood (10 μl) was collected at indicated time points from the tail vein for quantification of adoptively transferred pmel T cells by flow cytometry. Mice were euthanized when the total tumor volume exceeded 2,000 mm^3^, as per IACUC guidelines.

#### Toxicity evaluation

For measuring the toxicity of cytokines in vivo, mice were weighed and video recorded individually for 30 second intervals at baseline and every 48 hours as indicated. Videos were processed using EthoVision Software to quantify mobility.

#### T cell biodistribution assays

For biodistribution assays, mice were injected using B16-F10 tumor cells as previously described. 2.4 x 10^6^ non-sorted, transduced T cells were resuspended in 50 µl PBS and adoptively transferred five days after tumor inoculation through retro-orbital intravenous injection. Where indicated, mice received treatment with cytokines: MSA-bound mouse orthogonal IL-2 (MSA-oIL2), MSA-bound mouse IL-9 (MSA-IL9) (5 x 10^4^ units per day, intraperitoneal) for five doses.

The following tissues were harvested and dissociated into single-cell suspension at the indicated timepoints post adoptive cell transfer: spleen, tumor, liver, lung, blood. Spleen tissues were weighed, disintegrated mechanically, and filtered through a 70-µM cell strainer. Red blood cells were lysed with eBioscience RBC Lysis Buffer (1 mL per spleen; Thermo Fisher Scientific) for 5 minutes at 4°C and washed with PBS. Dissociated single-cell suspension splenocytes were strained in 70 µM-filters. Tumors were removed from the mouse, weighed, and minced until size was 2-4 mm. Minced tumors were added to tumor dissociation enzymes (Miltenyi Biotec) and dissociated using the gentleMACS dissociator for 40 minutes at 37°C, according to manufacturer’s protocols. Livers were removed from the mouse and weighed. Livers were subsequently added to liver dissociation enzymes (Miltenyi Biotec) and dissociated using the gentleMACS dissociator for 30 minutes at 37°C, according to manufacturer’s protocols. Lungs were removed from the mouse, perfused with PBS to remove residual blood, and weighed. Lungs were then separated into singe lobes and rinsed with PBS. Single lobes were subsequently added to lung dissociation enzymes (Miltenyi Biotec) and dissociated using the gentleMACS dissociator for 30 minutes at 37°C, according to manufacturer’s protocols. All single-cell suspensions of dissociated tissues were strained in 70 µM-filters. Peripheral blood (10 µl) was obtained through tail-vein sampling prior to collection of other organs. Red blood cells were lysed using ACK Lysing Buffer (Thermo Fisher Scientific) for 15 minutes at 4°C. After red blood cell lysis, cells were washed with PBS.

#### Mouse studies of human T cells in immunodeficient NSG mice

Nalm6-FFluc leukemia cells were cultured in complete RPMI media (10% fetal bovine serum, 100 U mL^-1^ penicillin, 100 μg mL^-1^ streptomycin, 2mM GlutaMAX, 10mM HEPES; Thermo Fisher Scientific). 1 x 10^6^ ml^-1^ Nalm6-FFluc cells in 200 µL sterile PBS were injected via tail vein into each mouse (n=5 per treatment group). On Day 3 post tumor inoculation, luciferin-based imaging was performed to confirm tumor engraftment. Upon confirmation, T cells were injected via tail vein at a dose of 1 x 10^6^ cells or 3 x 10^6^ cells in 200 µL (depending on the construct used) per mouse. Beginning on Day 3 as well, mice received a dose of 25,000 I.U. of MSA-hIL-9 in 100 µL of sterile PBS intraperitoneally. Mice were injected with cytokine every other day for a total of 10 doses.

Luciferase-based imaging was performed twice per week for at least three weeks to assess tumor growth. Specifically, mice were anaesthetized with isoflurane and 200 uL of 13mg mL^-1^ D-luciferin was injected i.p. per mouse, incubated for 4 minutes, and imaged for 30 seconds with heavy binning or automatic exposure detection. Mice were closely monitored and sacrificed upon reaching any one of the morbidity criteria, as outlined under IACUC guidelines (tumor flux exceeding 1 x 10^11^ p/s, rear leg paralysis, or other conditions that necessitated euthanasia). 143B-FFluc osteosarcoma cells were cultured in DMEM-c media. The right leg of each mouse was shaved, and width and depth were measured prior to tumor inoculation. 1 x 10^6^ 143B-FFluc cells in 100 µL sterile PBS were injected intramuscularly into the right leg of each mouse (n=5 per treatment group). On Day 4 post tumor inoculation, luciferin-based imaging was performed to confirm tumor engraftment. Upon confirmation, T cells were injected via tail vein at a dose of 7 x 10^6^ cells in 200 µL per mouse. Beginning on Day 4 as well, mice received a dose of 25,000 I.U. of MSA-hIL-9 intraperitoneally. Mice were injected with cytokine every over day for a total of 10 doses. Their right leg was measured three times a week. Mice were closely monitored and sacrificed upon reaching any one of the morbidity criteria, as outlined under IACUC guidelines (tumor size, inability to move their right leg, or other conditions that necessitated euthanasia).

### Coculture and cytotoxicity assays

Day 10 transduced human T cells and tumor cells were resuspended at 1 x 10^6^ cells ml^-1^ in RPMI-c with either no cytokine, 1 nM recombinant human IL-2, or 10 nM recombinant human IL-9 (Peprotech). T cells were serially diluted to achieve specified effector:target ratio and triplicates were cocultured with 5 x 10^4^ cells ml^-1^ (50uL) tumor cells. 50 uL of media was added to bring the total volume of each well to 200 uL. The plate was imaged in the Incucyte Live-Cell Analysis System (Sartorius) for 72 hours, and killing was quantified by measuring the Total Green Object Integrated Intensity metric of GFP+ tumor cells. Cytotoxicity index was calculated by normalizing to the fluorescence intensity at the first time point for each triplicate. Data were analyzed in Graphpad Prism. For repetitive killing assays, additional tumor cells were added in the same effector:target ratio as originally plated once tumor killing plateaued, while maintaining cytokine concentrations. In vitro proliferation assays were performed in triplicate in 96-well round-bottom plates. T cells, using AIM-V supplemented with 10% fetal bovine serum, 100 U/mL penicillin, 100 μg/mL streptomycin, 2mM GlutaMax, 10mM HEPES, 10 nM recombinant human IL-9 (Thermo Fisher Scientific, Peprotech). T cells were counted and percent live YFP+ cells were assessed by flow cytometry every 48 hours to calculate fold expansion, then re-plated in fresh media with cytokine at their original concentration. Flow cytometry data were analyzed using FlowJo and graphs were generated in GraphPad Prism.

### Phosphoflow signaling assay

Actively growing mouse or human primary T cells were rested at a concentration of 2 x 10^6^ cells mL^-1^ in T cell medium (either RPMI-c or complete AIM V for mouse and human, respectively) lacking IL-2 for 12 − 18 h before signaling assays. Cells were plated in a 96-well round bottom plate in T cell medium with 2% FBS for two hours. Cells were stimulated by addition of recombinant cytokines for 20 minutes at 37°C, and the reaction was terminated by fixation with 2.1% paraformaldehyde (BD Cytofix) for 30 minutes at 37°C. Cells were washed and permeabilized with ice-cold methanol (BD Phosflow Perm Buffer III) for 30 minutes on ice or stored at −20°C overnight. Cells were washed with staining buffer (eBioscience) before staining with pSTAT antibodies for 30 min to 1 h at 4°C in the dark. Cells were washed and analyzed on a CytoFlex (Beckman Coulter) or Novocyte Quanteon/Penteon (Agilent Technologies). Data represent the median fluorescence intensity (MFI), and points were fit to a log(agonist) versus dose–response model using Prism 10.2.3 (GraphPad). For the gating strategy, see Supplementary Figure 3E.

### Tandem mass tag (TMT)-based global phospho-proteomics

C57BL/6-derived mouse T cells were activated and transduced as previously described. Transduced T cells were sorted (2.5 x 10^7^ per replicate) based on expression of YFP^+^ using an Aria II cell sorter (BD Biosciences) and stimulated for 20 minutes at 37°C using the respective cytokines: rmIL-2 and rmIL-9 in T cell medium. Stimulated T cells were washed twice with PBS. Supernatant were removed, pellet was snap-frozen in liquid nitrogen and stored at −80°C before downstream analyses at the IDeA National Resource for Quantiative Proteomics.

Total protein from each sample was reduced, alkylated, and purified by chloroform/methanol extraction prior to digestion with sequencing grade trypsin and LysC (Promega). The resulting peptides were labeled using a tandem mass tag 10-plex isobaric label reagent set (Thermo), combined into two multiplex sample groups, then enriched using High-Select TiO2 and Fe-NTA phosphopeptide enrichment kits (Thermo) following the manufacturer’s instructions. Both enriched and un-enriched labeled peptides were separated into 46 fractions on a 100 x 1.0 mm Acquity BEH C18 column (Waters) using an UltiMate 3000 UHPLC system (Thermo) with a 50 min gradient from 99:1 to 60:40 buffer A:B ratio under basic pH conditions, then consolidated into 18 super-fractions for the enriched and un-enriched sample sets (Buffer A = 0.1% formic acid, 0.5% acetonitrile; Buffer B = 0.1% formic acid, 99.9% acetonitrile). Both buffers were adjusted to pH 10 with ammonium hydroxide for offline separation. Each super-fraction was then further separated by reverse phase XSelect CSH C18 2.5 um resin (Waters) on an in-line 150 x 0.075 mm column using an UltiMate 3000 RSLCnano system (Thermo). Peptides were eluted using a 75 min gradient from 98:2 to 60:40 buffer A:B ratio. Eluted peptides were ionized by electrospray (2.4 kV) followed by mass spectrometric analysis on an Orbitrap Eclipse Tribrid mass spectrometer (Thermo) using multi-notch MS3 parameters. MS data were acquired using the FTMS analyzer in top-speed profile mode at a resolution of 120,000 over a range of 375 to 1500 m/z. Following CID activation with normalized collision energy of 31.0, MS/MS data were acquired using the ion trap analyzer in centroid mode and normal mass range. Using synchronous precursor selection, up to 10 MS/MS precursors were selected for HCD activation with normalized collision energy of 55.0, followed by acquisition of MS3 reporter ion data using the FTMS analyzer in profile mode at a resolution of 50,000 over a range of 100-500 m/z. Proteins were identified and reporter ions quantified by searching the UniprotKB database using MaxQuant (Max Planck Institute) with a parent ion tolerance of 3 ppm, a fragment ion tolerance of 0.5 Da, a reporter ion tolerance of 0.001 Da, trypsin/P enzyme with 2 missed cleavages, variable modifications including oxidation on M, Acetyl on Protein N-term, and phosphorylation on STY, and fixed modification of Carbamidomethyl on C. Protein identifications were accepted if they could be established with less than 1.0% false discovery. Proteins identified only by modified peptides were removed. Protein probabilities were assigned by the Protein Prophet algorithm [*Anal. Chem.* **75:** 4646-58 (2003)]. TMT MS3 reporter ion intensity values are analyzed for changes in total protein using the unenriched lysate sample. Phospho(STY) modifications were identified using the samples enriched for phosphorylated peptides. The enriched and un-enriched samples are multiplexed using two TMT10-plex batches, one for the enriched and one for the un-enriched samples.

Following data acquisition and database search, the MS3 reporter ion intensities were normalized using ProteiNorm (Graw et al). The data was normalized using VSN (Huber et al) and analyzed using ProteoViz to perform statistical analysis using Linear Models for Microarray Data (limma) with empirical Bayes (eBayes) smoothing to the standard errors (Storey et al, Ritchie et al). A similar approach is used for differential analysis of the phosphopeptides, with the addition of a few steps. The phosphosites were filtered to retain only peptides with a localization probability > 75%, filter peptides with zero values, and log2 transformed. Limma was also used for differential analysis. Proteins and phosphopeptides with an FDR-adjusted p-value < 0.05 and an absolute fold change > 2 were considered significant.

### Mouse T cell in vitro expansion, proliferation (CellTrace) and apoptosis/cell death assays

Actively growing mouse T cells were rested for 24 hours post-transduction in T cell medium with 100 U ml^-1^ rmIL-2. On day of assay, T cells were washed with T cell medium lacking cytokine. IL-9R transduced T cell proliferation was assessed by seeding 100,000 cells per well in a round bottom 96-well plate in the presence of equipotent doses of recombinant murine IL-2 (rmIL-2), rmIL-9, or rmIL-12. T cells were fed with fresh media and cytokine 48 hours after seeding, and every subsequent 48 hours until the end of the assay. Counts were also acquired on the day of replenishment by staining an aliquot of cells with Acridine Orange/Propidium Iodide (AO/PI, The DeNovix Company) and analyzing on a cell counter (The DeNovix Company). An aliquot of cells were also stained with Zombie Violet Live Dead (Biolegend) and analyzed for YFP^+^ on the NovoCyte Penteon (Agilent Technologies). For CellTrace assays, resuspension of dye was followed as recommended by the manufacturer. In short, cells were resuspended at 1 x 10^6^ cells/mL and labeled with CellTrace Violet (Thermo Fisher) at the following ratio :1 × 10^6^ cells per 1 mL of 10 µM Cell Trace. Cell mix was subsequently incubated at 37°C for 20 minutes with occasional vortexing. Cells were subsequently diluted 5x in T cell medium and incubated at 37°C for 5 minutes. Cells were then washed and resuspended at a concentration of 5 x 10^5^ cells/mL and plated in a 96-well plate with the respective cytokine. To measure apoptosis, cells were removed every 48 hours post cytokine treatment. Cells were subsequently resuspended at 1 x 10^6^ cells/mL and stained with Zombiet Violet (BioLegend) and Annexin V (Thermo Fisher) in Annexin Binding buffer (Biolegend) at 25°C for 15 minutes. After the incubation, cells were diluted with 2x Annexin Binding buffer, placed on ice, and immediately analyzed on the Novocyte Quanteon (Agilent Technologies).

### Immunophenotyping by Flow Cytometry

For in vitro immunophenotyping of transduced T cells, non-sorted T cells were plated with equipotent doses of rmIL-2 or rmIL-9 in triplicates. After 24 h or 48 h, T cells were collected and surface stained. Immediately after staining, cells were fixed in 1.6% paraformaldehyde (PFA) for 15 minutes at 25°C and washed before analysis on the LSR II (BD Biosciences). For in vivo assessments, peripheral blood was obtained through tail-vein sampling at indicated timepoints. Red blood cells were lysed using ACK Lysing Buffer (Thermo Fisher Scientific) for 10 minutes at 4°C. After red blood cell lysis, cells were washed with PBS and stained. Cells were fixed in 1.6% paraformaldehyde (PFA) for 15 minutes at 25°C and washed before analysis on NovoCyte Penteon (Agilent Technologies).

### ELISA

Serum IFNγ ELISA was conducted using the Mouse IFNγ ELISA MAX kit (Biolegend) using the manufacturer’s recommendations. In short, the day prior to running the ELISA, the Capture Antibody was diluted in Coating Buffer and 100 µL of this capture solution to all relevant wells of a 96-well plate. The plate was sealed and incubated overnight at 4°C. On the day of running the ELISA, the plate was washed 4 times with 300 µL of Wash Buffer per well to remove capture solution. After the wash was complete, the plate was then blocked with 200 µL of Assay Diluent per well, and then incubated at 25°C for 1 hour on a plate shaker (500 rpm). Following the blocking, the plate was washed 4 times with Wash Buffer, and then 100 µL of standard dilutions and samples were added in duplicates. The plate was then incubated at 25°C for 2 hours with shaking. After the incubation, the plate was washed 4 times as previously described and 100 µL of diluted Detection Antibody solution was added to each well and incubated at 25°C for 1 hour with shaking. Following the detection incubation, the plate was washed 4 times and add 100 µL of diluted Avidin-HRP solution was added and incubated at 25°C for 30 minutes with shaking. After the incubation, the plate was washed 5 times with Wash Buffer, with the buffer soaking in each well for 30 seconds per wash to minimize background. After the washes were complete, 100 µL of TMB Substrate Solution was added and incubated at 25°C in the dark until the wells turn blue. The reaction was stopped by adding 100 µL of Stop Solution to each well and then immediately read on a fluorescent microplate reader (Thermo Fisher Scientific) (absorbance read at 450 nm).

### Cytokine dose calculations

For in vivo experiments, we opted to utilize the activity of the cytokine (international units; I.U.) as opposed to concentration (µg). We enacted the following conversion factors for the following cytokines: For MSA-mIL2, MSA-hIL2 or MSA-oIL2 (3A10), we considered 1 µg of protein to equal 3280 I.U. For MSA-mIL9, we considered 1 µg to equal 1118 I.U. For MSA-hIL9, we considered 1 µg MSA-IL9 equal to 1428 I.U.

### Analysis of in-house single cell RNA-sequencing data

Single cell RNA-sequencing of sorted live thy1.1+YFP+ pmel T cells was performed using the 10X Genomics platform using 10x Genomics reagents according to the manufacturer’s instructions. The 3′ Gene Expression libraries were sequenced using a NovaSeq 6000 with a sequencing depth of 120M paired reads (240M total reads) per sample to achieve >20K reads per cell.

#### Data pre-processing

The FASTQ files were processed using the cellranger count pipeline [CellRanger version 7.0.1 (10x Genomics)] using the mm10 mouse reference genome. Raw gene expression matrices were constructed into a seurat object and imported into R software using Seurat (version 5). To filter out low-quality cells, we removed cells that had either a low or high number of detected genes and cells that had more than 10-15% of mitochondrial UMI counts in the scRNA-seq data and remaining 6706 cells were used for downstream analysis. To normalize the gene expression levels, we utilized the LogNormalize method implemented in Seurat. We performed principal component analysis (PCA) to reduce the dimensionality of scRNA-seq data.

#### Cell clustering and cell type annotation

Top 30 principal components (PCs) were selected to construct the UMAP embeddings. The FindClusters function was used to identify cell clusters. Total of 10 major cell clusters were obtained, and cluster specific markers were identified using “FindAllMarkers” or “FindMarkers” functions in Seurat. Only positive markers expressed in at least 1% of cells were considered. The nonparametric Wilcoxon rank-sum test was used to obtain the *p* value for comparisons, and the adjusted *p* value based on Bonferroni correction was calculated. Genes with adjusted p < 0.05 were considered as enriched in a particular cluster.

#### Trajectory analysis

To simulate and understand the differentiation trajectories between cells, we used “monocle3” package (version 1.3.7). The T cell clusters and differential gene expression analysis with Seurat were extracted from the scRNA-seq data and used to construct the monocle object. Pseudotime trajectories were constructed with default parameters and visualized with the plot_cells() function. Cells from the “Tscm-like (Tcf7)” cluster were used as the reference starting point (root nodes) from which potential differentiation pathways resonate. The identified paths were mapped on to the UMAP projection for visualization. Ridge plots shows the pseudotime scores across treatment groups or clusters.

#### Transcription factor analysis in scRNA-seq data

Top regulons (gene sets regulated by the same transcription factor) and their activity were inferred and evaluated using the SCENIC pipeline (version 1.3.1), thereby defining the top regulon activity in every cell. The input files consisted of the expression matrix and metadata. Then, the co-expression network was calculated by *GRNBoost2*, and the regulons were identified by *RcisTarget*. Next, the regulon activity for each cell was scored by *AUCell*. Scaled expression of regulons were used for the plotting.

#### Gene set signature scores for scRNA-seq data

“AddModuleScore” function in Seurat with default parameters was used to score sets of genes in the single cell data. We calculated the average expression levels of each gene set on a single cell level, subtracted by the aggregated expression of control gene sets. All analyzed features are binned based on averaged expression, and the control features are randomly selected from each bin. The scored expression was used for plotting.

#### Statistical analysis

Statistical comparisons between cell clusters, treatment conditions and regulon activity were performed using R. Kruskal-Wallis test to compare the medians of groups. *P* < 0.05 was considered statistically significant.

### Analysis of in-house bulk RNA-sequencing data

Libraries were prepared using KAPA mRNA HyperPrep Kit from total RNA. Paired-end sequencing was performed on a Novaseq 6000/X Plus (40M reads per sample).

#### Data pre-processing

Read alignment was performed using STAR aligner on GRCm38 mouse reference transcriptome and transcript level quantification using Salmon. Quality control of the paired-end bulk RNA-seq data was performed using the FastQC program. Adapter trimming was carried out by Cutadapt method. Initial quality control of the count data was carried out by PCA and sample-sample distance after variance stabilizing normalization. DESeq2 was used to perform the differential gene expression analysis on samples.

#### Transcription factor analysis in bulk RNA-seq data

To infer TF activity in bulk RNA-seq data we ran the Univariate Linear Model (ulm) method using the decoupleR package. For each sample in our dataset (mat) and each TF in our network (net), we fit a linear model that predicts the observed gene expression based solely on TF-Gene interaction weights. Once fitted, the obtained t-value of the slope is the score. If it is positive, we interpret that the TF is active and if it is negative we interpret that it is inactive.

#### Gene set scoring for bulk RNA-seq data

We calculated the gene set score of the bulk RNA-seq data based on the average normalized expression of the genes. From this, the enrichment score of randomly selected control genes with similar expression levels are subtracted. These control gene sets are defined by first binning all genes into bins (n= 24) of aggregate expression level and then, for each gene in the gene set of interest, 100 genes from the same expression bin as that gene are randomly selected. The score is then set to range from zero, meaning no enrichment compared to random sets of genes with similar expression, to one, reflecting the highest average expression of all genes within the gene set of interest. This gene set score is calculated per sample for bulk RNA-seq samples.

#### Functional Enrichment Analysis

Functional geneset enrichment analysis was performed using fGSEA package in R to infer enriched biological pathways in the treatment groups.

#### Linking bulk RNA-seq with phosphoflow data to generate pSTAT gene modules

We utilized a set of samples with matched RNA-seq and phosphoflow data. The RNA-seq data was obtained from in vitro experiments, while the phosphoflow data consisted of MFI measurements at maximum stimulation (Emax) for phosphorylated STAT proteins (pSTAT1, pSTAT3, pSTAT4, and pSTAT5). These two datasets were merged to create a combined matrix for subsequent analysis. To investigate the relationship between gene expression and STAT phosphorylation, we computed Pearson correlation coefficients between the RNA-seq counts and each pSTAT measurement. This analysis was performed using the cor.test function from the R stats package. The correlation test was conducted for each gene against each pSTAT, resulting in a comprehensive correlation matrix. For each pSTAT, we selected the top 100 genes exhibiting the highest absolute correlation coefficients. These genes were then visualized using scatter plots and the coefficient of determination (R²) was calculated for each pSTAT-gene set relationship to quantify the proportion of variance in pSTAT levels explained by gene expression. We created gene modules for each pSTAT using the top 100 correlated genes identified in the previous step and applying the AddModuleScore function in Seurat. These gene modules were then projected onto our single-cell RNA sequencing (scRNA-seq) dataset as violin and feature plots.

### Analysis of publicly available single cell and bulk RNA-seq datasets

#### PBMC and lamina propria single cell RNA-seq datasets

The raw expression data was obtained from the Single Cell Portal (Broad Institute) for Human PBMC (“Single Cell Comparison: PBMC data”)^12^ and Lamina Propria (“ICA: Ileum Lamina Propria Immunocytes”)^13^. Raw gene expression matrices were constructed into a Seurat object and imported into R software using Seurat (version 5). Data processing was performed as described in sections above. t-Distributed stochastic neighbor embedding (t-SNE) in Seurat was used for data visualization. Azimuth (version 0.5.0) in Seurat was used to annotate the clusters obtained at resolution 0.2 (PBMC) and 0.5 (lamina propria). To specifically visualize the expression of γ_c_ cytokines and their receptors, we plotted feature plots and dot plots to examine the expression across different cell types in PBMC and Lamina Propria.

#### Human Protein Atlas bulk RNA-sequencing data

RNA GTEx (genotype tissue expression) tissue gene data was downloaded from The Human Protein Atlas^61^ (v19.proteinatlas.org/about/download). Transcript expression levels summarized per gene in 36 tissues based on RNA-seq. The tab-separated file includes Ensembl gene identifier (“Gene”), analysed sample (“Tissue”), transcripts per million (“TPM”), protein-transcripts per million (“pTPM”) and normalized expression (“NX”). The data was obtained from GTEx and is based on The Human Protein Atlas version 19.3 and Ensembl version 92.38. The expression values for *IL2RB, IL4R, IL7R, IL9R*, and *IL21R* across different tissues were used for visualization. We defined a threshold of 0.7, below which expression is not detected or negligible.

#### Single cell RNA-seq data of mouse lymph nodes in response to cytokines

Data was obtained from previously published work on the single cell transcriptomic responses of mouse immune cells (from lymph nodes) in response to a large panel of different cytokines^17^. Data were downloaded as single cell data objects that were reanalyzed to quantify the overall magnitude of transcriptomic responses of each cytokine across different cell types. We created a dot plot with two metrics: the number of differentially expressed genes (DEGs) and the magnitude of change (based on Euclidean distance) across the entire transcriptome. The number of DEGs was the total number of genes in each cytokine signature. The overall magnitude of cytokine-induced differential expression was computed as the Euclidean distance between the centroid vectors of cytokine-treated cells and PBS-treated cells. A distinct color ramp was used for each cell type to emphasize that cell types have different properties (e.g., different numbers of genes expressed on average) and were independently analyzed. Cytokine–cell type combinations with five or more cells sampled were included in this analysis.

#### Single cell RNA-seq data from human CAR T cells sorted from patients

We utilized publicly available data from a single-cell RNA sequencing analysis of sorted CAR T cell populations at various timepoints after patient administration^20^. Across all patients and time points, the authors sequenced 66,042 post-infusion CAR T cells, with an average of 11,549 cells per patient (SD = 7,335) and 20,532 cells per time point (SD = 37,898). Notably, the month 6 post-infusion time point only consists of 7 CAR T cells. We downloaded Seurat objects containing different samples at various time points, fetched the expression of relevant cytokine receptors across different cell types and generated dot plots of the data.

### Soft tissue sarcoma single-cell RNA-seq samples and related analysis

Previously published synovial sarcoma 10x Genomics and SMART-seq2 scRNA-seq data were downloaded from GEO (GSE131309)^62^. The undifferentiated pleomorphic sarcoma (UPS) and myxofibrosarcoma (MFS) samples were obtained GEO (GSE212527)^63^. Log-normalization was performed independently for each sample using the Seurat function NormalizeData. After this step, the Seurat objects for each independent sample were merged preserving the individual normalization of the data. Sample integration and clustering were performed separately for each histotype, accounting for batch effects and differences in sequencing technologies. The top 2000 highly variable genes of the merged and normalized expression matrices were identified using the function FindVariableFeatures, and then centered and scaled using the ScaleData function. Principal component analysis was then run using the top 2000 highly variable genes previously identified. To integrate multiple samples, the harmony R package*(3)* (version 0.1.0) was employed for batch correction. Harmony was ran using both the sample of origin and the sequencing technology variables as arguments of “group.by.vars”, and based on the first 25 PCA dimensions previously identified. UMAP embeddings were then obtained using the first 25 Harmony dimensions, and clusters were obtained by calculating the k-nearest neighbors (k-NN) and a shared nearest-neighbor graph using the Louvain algorithm implemented in the FindClusters function of the Seurat package with a resolution of 1.0. Endothelial cells, myeloid cells, lymphoid cells, and CAFs were annotated using the differentially expressed genes discovered with the FindMarkers function in Seurat, which was run with the default parameters except for only.pos = T and min.pct = 0.25. Canonical cell identity markers obtained from the literature were used for cell type annotations. A small number of cells could not be uniquely assigned to a specific cell identity and were removed from further downstream analysis. UMAP plots were generated with Seurat and scCustomize *(4)*. Density plots were generated with the R package Nebulosa *(5)*.

### Statistical analysis

Statistical analysis was performed using GraphPad Prism 9 (GraphPad software), except where indicated. All values and error bars are shown as mean ± SEM. Comparisons of two groups were performed by using two-tailed unpaired Student’s t test. Comparisons of multiple groups were performed by using one-way analysis of variance (ANOVA) with Tukey’s multiple-comparisons test unless otherwise indicated. Experiments that involved repeated measures over a time course, such as tumor growth were performed by using two-way ANOVA with Tukey’s multiple-comparisons post-test. Survival data were analyzed using the Log-rank (Mantel-Cox) test.

## Ethics statement

Experimental procedures in mouse studies were approved by the Institutional Animal Care and Use Committee (IACUC) at the Stanford University (animal protocol ID 34340 and 33698) and performed in accordance with the guidelines from the animal facility of Stanford University. Primary T lymphocytes from healthy donors were provided by the Stanford Blood Center. Ethical approval pertaining to T cell donors was obtained by the Stanford Blood Center.

## Data availability

The raw and processed scRNA-seq data generated in this study will be deposited for public availability. The R code used to analyze the scRNA-seq data will be made publicly available before publication. Analysis details are provided in the Methods section. Any additional information required to reanalyze the data reported in this paper is available from the lead contact upon request.

## Acknowledgements

This work was supported by the NIH R37CA273074 (A.K.), NIH K08CA245181 (A.K.), the Damon Runyon Cancer Research Foundation (A.K.), NIH-RO1AI51321 (K.C.G.), Ludwig Institute (K.C.G.), and the Parker Institute of Cancer Immunotherapy (PICI) (A.K., K.C.G); K.C.G. is an investigator with the Howard Hughes Medical Institute. Cell sorting/flow cytometry analysis for this project was done on instruments in the Stanford Shared FACS Facility (RRID: SCR_017788). Additionally, data was collected on an instrument in the Shared FACS Facility obtained using NIH S10 Shared Instrument Grant (1S10OD026831-01). Phospho-TMT analysis was conducted by the IDeA National Resource for Quantitative Proteomics and NIH/NIGMS grant R24GM137786. We thank Crystal Mackall and Louai Labanieh for sharing cell lines and CAR constructs and for valuable scientific feedback.

## Author contributions

A.K. conceived the study, acquired funding, designed and supervised experiments, and wrote the manuscript. S.L., H.J., K.K., L.W.R., O.L., L.S. and H.O. conducted experiments and assisted with experimental design. L.S. conducted an original phosphotyrosine mutant screen with o9R that served as a basis for the IL-9R screen. S.L., H.J., K.K., L.W.R., A.P. and O.L. analyzed the data. S.L., K.K., L.W.R., and A.P. contributed to manuscript writing. A.P. and A.K. analyzed bulk RNA-seq and scRNA-seq data; S.T. and E.J.M. assisted with scRNA-seq analysis. K.C.G. assisted with experimental design and interpretation and supervised protein production. P.T. assisted with design of the IL-9R^3x^ and IL-9R^5x^ mutants. D.W. assisted with protein production. K.J. assisted with protein structure prediction. C.S.S. and G.M.C. provided RNA-sequencing datasets. All authors revised and edited the manuscript.

## Supplemental Information

Figures S1–S7 (appended to manuscript).

Table S1. Differentially phosphorylated proteins, related to Figure 3C.

Table S2. Differential gene expression and regulon analysis, related to Figure 6.

Table S3. Correlations between STAT phosphorylation and gene expression, related to Figure 6H-K.

**Figure S1.**
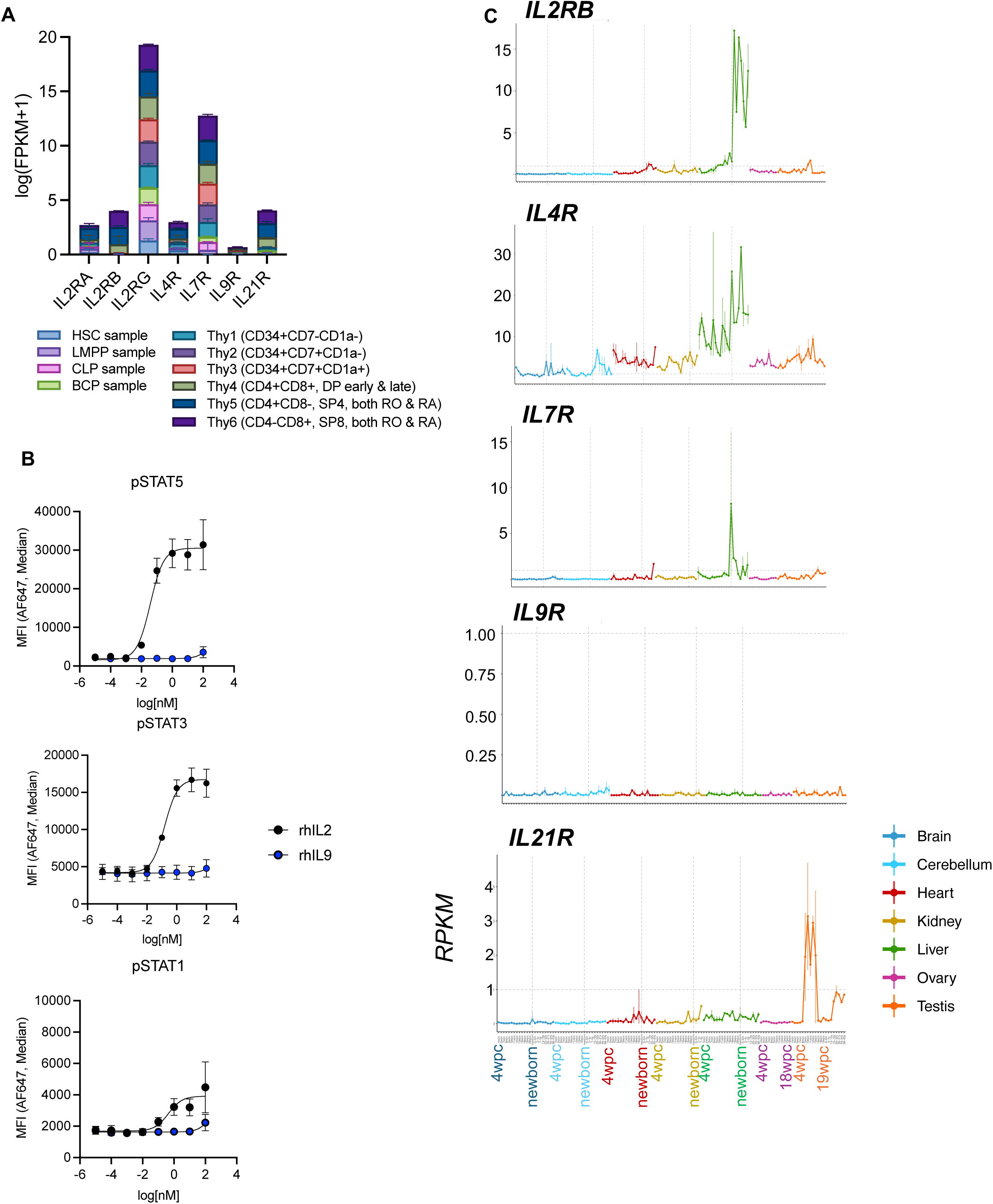
Nomination of IL-9 as a naturally orthogonal cytokine, related to Figure 1. (A) Expression [log(FPKM+1)] of receptors for γ_c_ cytokines from publicly available RNA sequencing data from sorted populations from normal bone marrow and thymus of human donors representing different stages of hematopoietic and T cell development^14^. (B) Dose-response curves of STAT1, STAT3, and STAT5 phosphorylation in isolated healthy donor human T cells (n=4 healthy donors) stimulated on Day 6 post activation with either recombinant human IL-2 or IL-9 for 20 minutes. (C) Expression of receptors for γ_c_ cytokines from publicly available RNA sequencing data of 22 stages of fetal and human development spanning 4 weeks post conception (4wpc) through 63 years of age. Plot is colored by tissue type. Vertical dashed lines indicate newborn stage for each tissue type. Horizontal dashed lines indicate threshold RPKM of 1.

**Figure S2.**
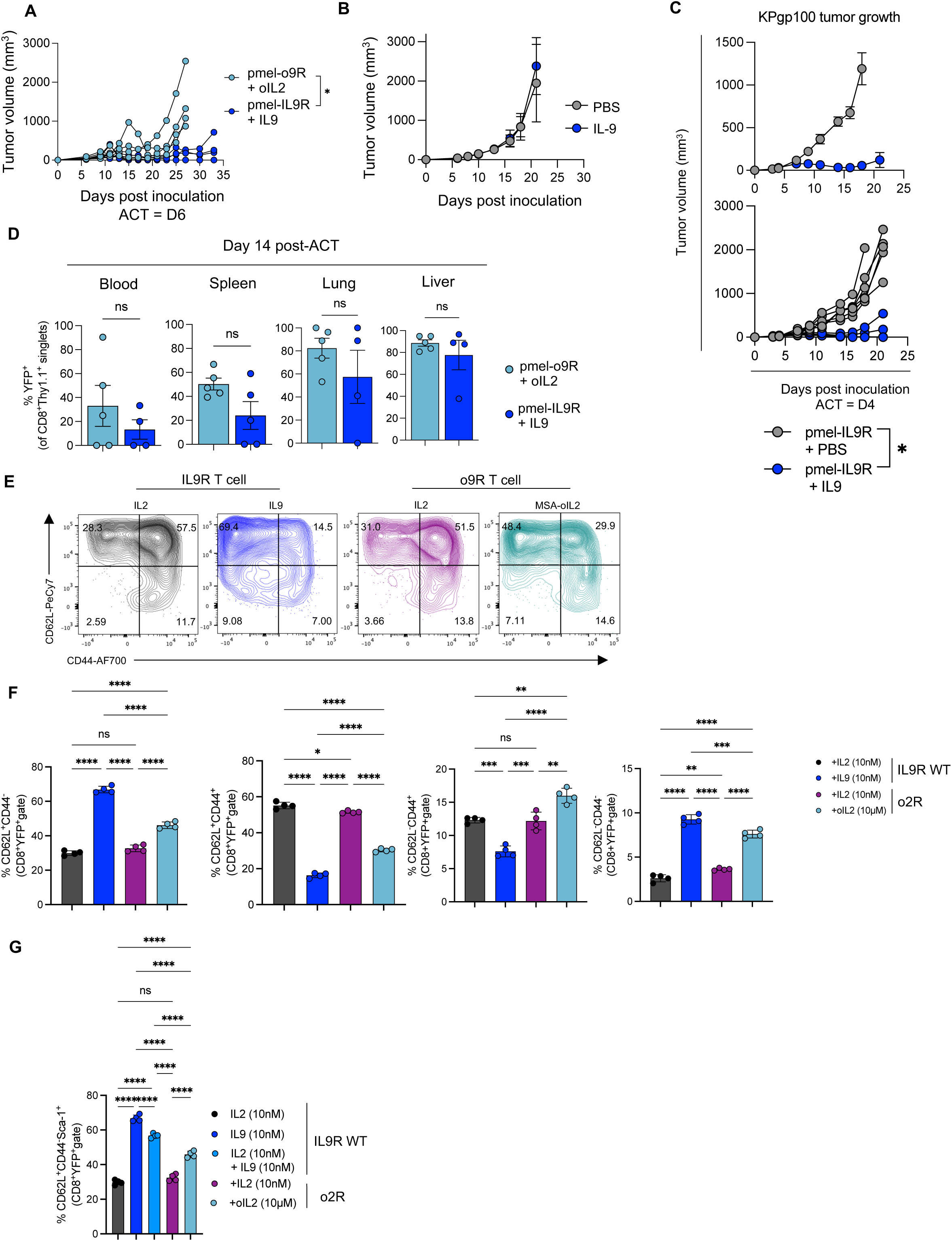
Anti-tumor and phenotypic effects of IL-9, related to Figure 2. (A) Tumor growth (from Figure 2F) with data shown for individual mice. (B) B16-F10 tumor growth (mean ± SEM, n=5 mice/group) in mice treated with either PBS or IL-9 (5×10^4^ IU, i.p/dose) starting five days after tumor inoculation and continuing every other day for five doses. (C) KP-gp100 tumor growth (mean ± SEM and individual mice, n=6 mice / group) after ACT with pmel T cells engineered with IL-9R (1.6×10^6^ transduced cells, i.v.), and cytokine treatment with PBS or IL-9, respectively (5×10^4^ IU i.p., every other day for 5 days starting with ACT). Data is representative of two independent experiments. (D) Quantification of transduced pmel T cells (Thy1.1^+^YFP^+^) across different tissues 14 days after ACT and five doses of MSA-oIL2 or MSA-mIL9 (n=5 mice/group). (E) Representative gating strategy of CD44 and CD62L phenotyping for IL-9R or o9R-engineered T cells 24h after stimulation with IL-2 (10nM), IL-9 (10nM), or MSA-oIL2 (10µM). See Figure 2K. (F) Related to Figure 2K-L. The percentage of naïve (CD62L+CD44-), central memory (CD62L+CD44+), effector (CD44+CD62L-) or double negative (CD62L-CD44-) T cells as bar plots. (G) Proportion of T_SCM_ cells (CD44-, CD62L+, Sca-1+) from (E) presented as bar plots. For all in vivo experiments, cytokines were tagged with mouse serum albumin (MSA) for half-life extension. *P < 0.05, **P < 0.005, ***P < 0.0005, ****P < 0.0001 (two-way ANOVA for B; Welch’s T-test for C; one-way ANOVA for D; two-way ANOVA for F-G).

**Figure S3.**
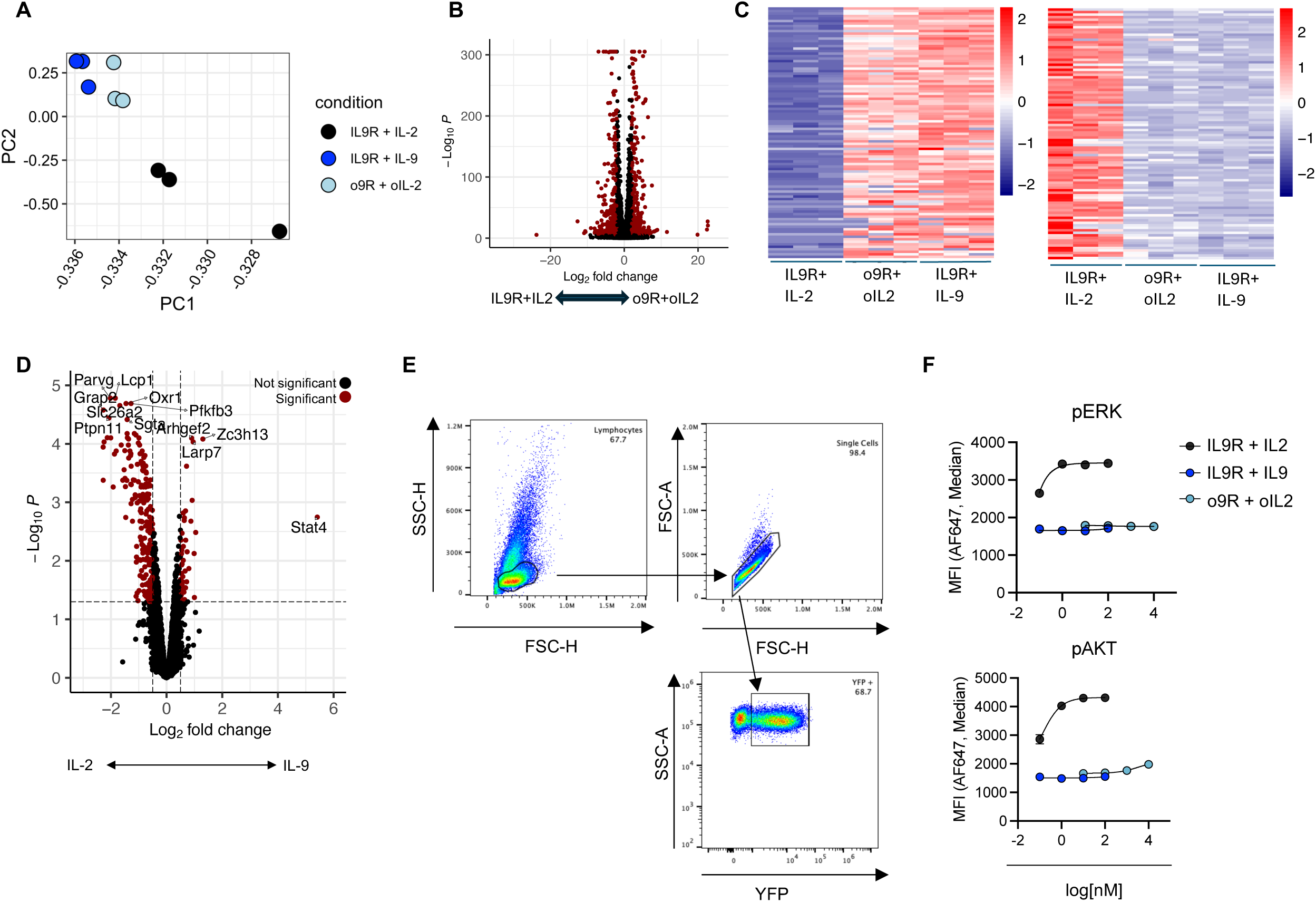
Signaling and transcriptomic effects of IL-9R engagement, related to Figure 3. (A) Principal component analysis (PC1 vs PC2) of RNA sequencing of C57BL/6 T cells transduced for 48h with IL-9R and treated with IL-9 (10 nM) or IL-2 (10 nM), or with o9R and treated with oIL2 (10 µM). Samples cluster by treatment. (B) Volcano plots depicting differential gene expression based on RNA-sequencing of C57BL/6 T cells transduced with IL-9R or o9R and treated with IL-9 (10 nM), IL-2 (10 nM) or oIL2 (10 µM) for 48 hours. Shown here is the comparisons between o9R T cells treated with oIL-2 and IL-9R T cells treated with IL-2. Significance (red) indicates adjusted p-value < 1 x 10^-5^. (C) Heatmap of top 100 differentially upregulated (left) and downregulated (right) transcripts in T cell expressing IL-9R treated with IL-9 versus IL-2 for 48h. Shown also are the o9R samples treated with oIL-2, which mimic the expression of the IL-9R + IL-9 samples. (D) Differentially phosphorylated proteins between C57BL/6 T cells transduced with IL-9R and stimulated with IL-2 (10nM) or IL-9 for 20’. Significance (red) indicates adjusted p < 0.05 and log_2_(fold change) ≥ 0.5. (E) Representative gating strategy for flow cytometry data for intracellular staining of phosphoproteins in engineered (YFP^+^) T cells. (F) Dose-response curves of ERK or AKT phosphorylation in IL-9R or o9R transduced pmel T cells (YFP^+^) and stimulated with either oIL-2, IL-2, or IL-9 for 20 minutes. Error bars represent min/max of technical duplicates. Data representative of two independent experiments.

**Figure S4.**
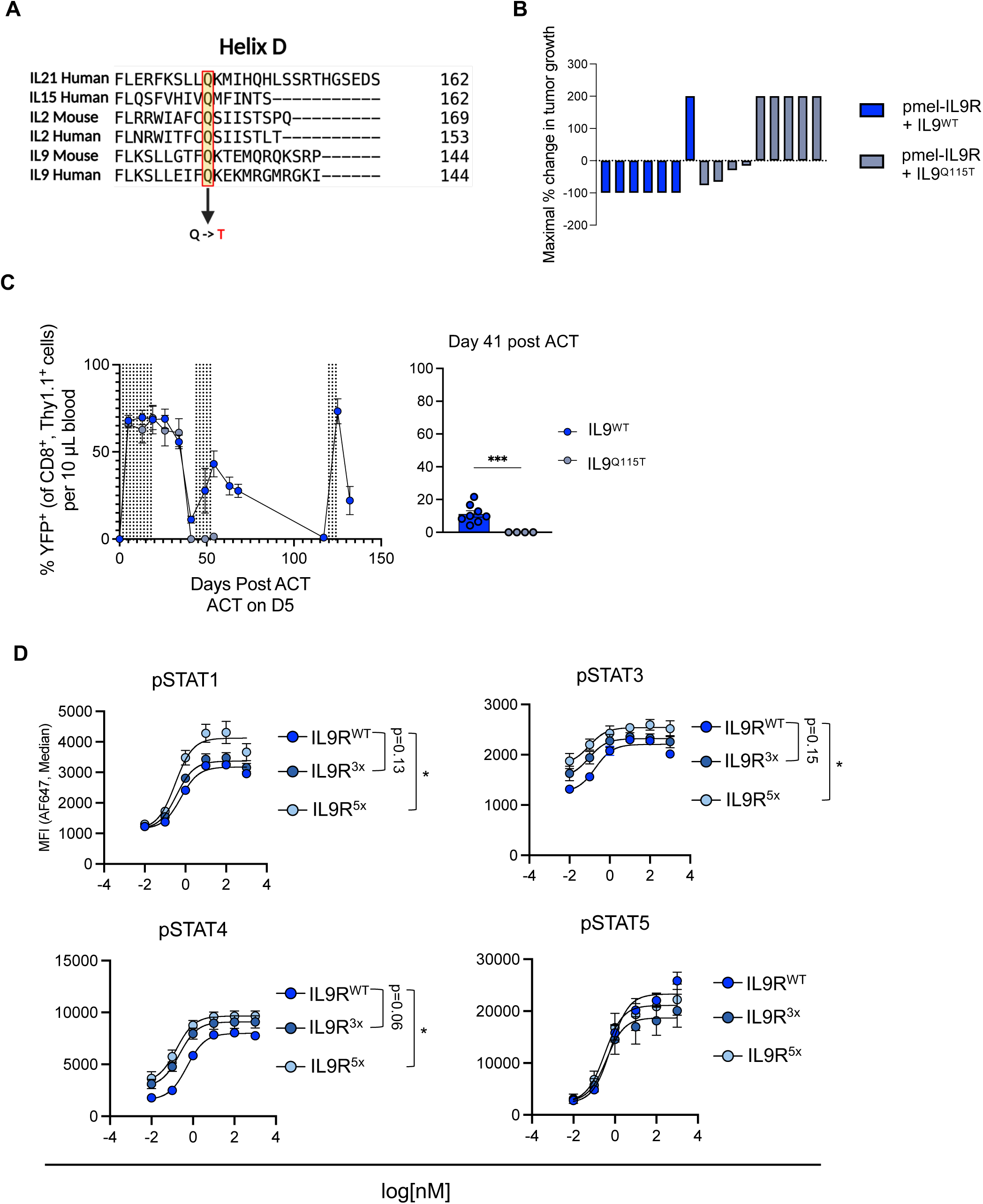
Structure-based attenuation or amplification of IL-9/IL-9R, related to Figure 4. (A) Sequence alignment between mouse and human IL-2, IL-9, IL-15, and IL-21 around the conserved glutamine in Helix D. (B) Waterfall plot of maximal tumor size reduction for experiment from Figure 4C, in which B16-F10 tumors in mice (n=7-9 mice/group) treated with IL-9R transduced pmel T cells and either IL-9^WT^ or IL-9^Q115T^. (C) Peripheral blood enrichment of IL-9R transduced pmel T cells over time in mice treated with ACT and either IL-9^WT^ or IL-9^Q115T^ cytokine (each cytokine dose indicated by a vertical dashed line). Enrichment was calculated as percentage of transduced (YFP+) pmel T cells as a proportion of all pmel T cells. See also Figure 4D. (D) Dose-response curves of phosphorylation of indicated STAT proteins in pmel T cells transduced with IL-9R^WT^, IL-9R^3x^, and IL-9R^5x^ and stimulated with IL-2 or IL-9 for 20 minutes. Error bars represent min/max of three biological replicates. For in vivo experiments, cytokines were tagged with mouse serum albumin (MSA) for half-life extension. ***P < 0.001 (Welch’s t-test for C, two-way ANOVA for D).

**Figure S5.**
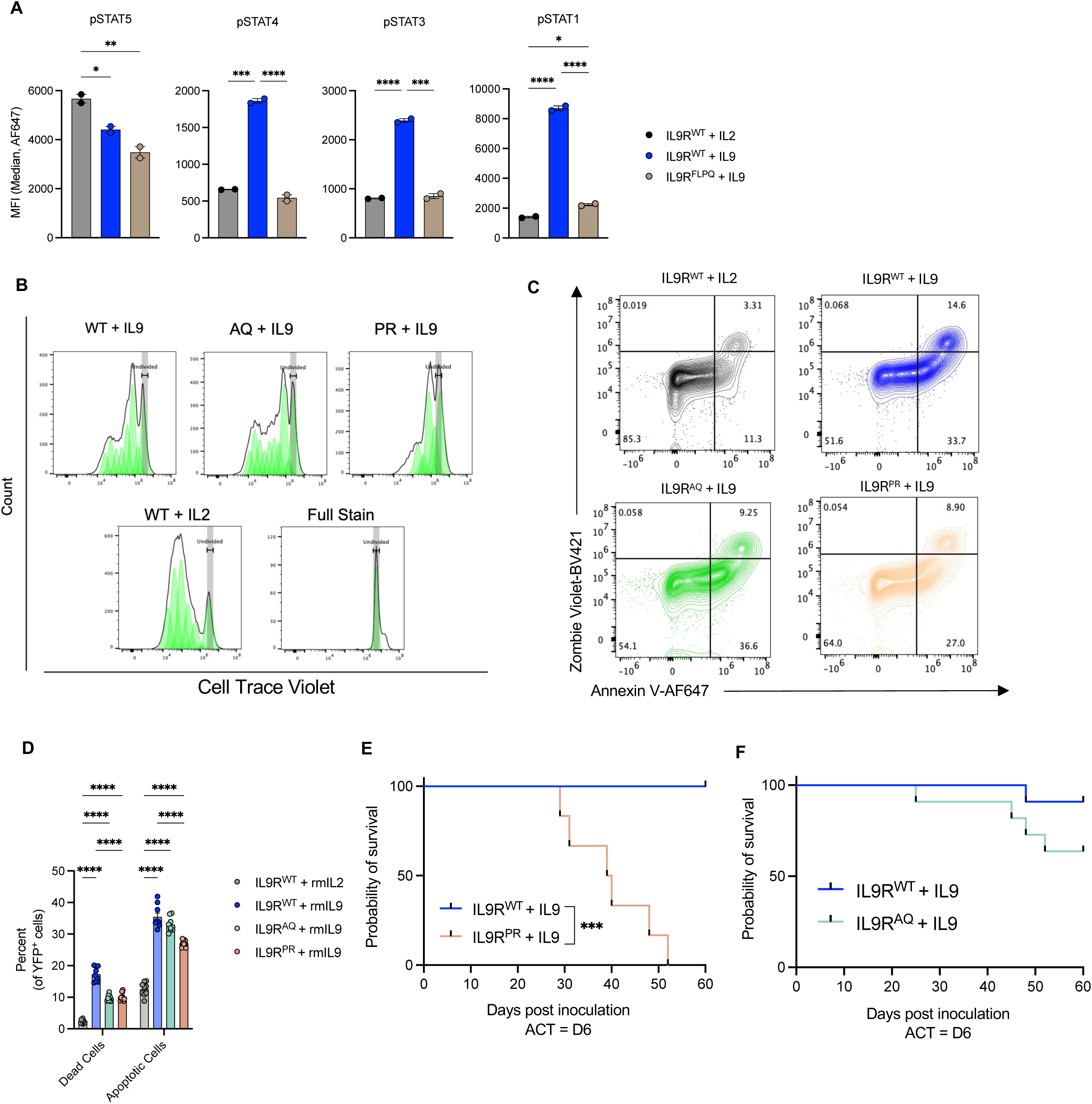
Proliferation of IL-9R and IL-9R intracellular domain mutants, related to Figure 5. (A) Phosphorylation of indicated STATs at E_max_ (10nM) for IL-9R^WT^, IL-9R^FLPQ^ transduced C57BL/6 T cells stimulated with IL-2 or IL-9 for 20 minutes. Bars represent mean ± SEM of technical replicates. Data is representative of two independent experiments. (B) Representative histograms of CellTrace Violet staining for the groups from Figure 5E. (C) Percentage of YFP^+^ cells undergoing apoptosis (Annexin V^+^, Zombie Violet^-^) or dead (Annexin V^+^, Zombie Violet^+^) shown as bar plots (each point represents the average of technical replicates for each biological replicate). (D) Related to Figure S5D. Percentage of YFP^+^ cells undergoing apoptosis (Annexin V^+^, L/D^-^) or dead (Annexin V^+^, L/D^+^) shown as bar plots (each point represents the average of technical replicates for each biological replicate). (E) Survival curves of mice from Figure 5G (F) Survival curves of mice from Figure 5H *P < 0.05, **P < 0.005, ***P < 0.001, ****P < 0.0001 (one-way ANOVA for A-B, two-way ANOVA for E, Mantel-Cox test for F-G).

**Figure S6.**
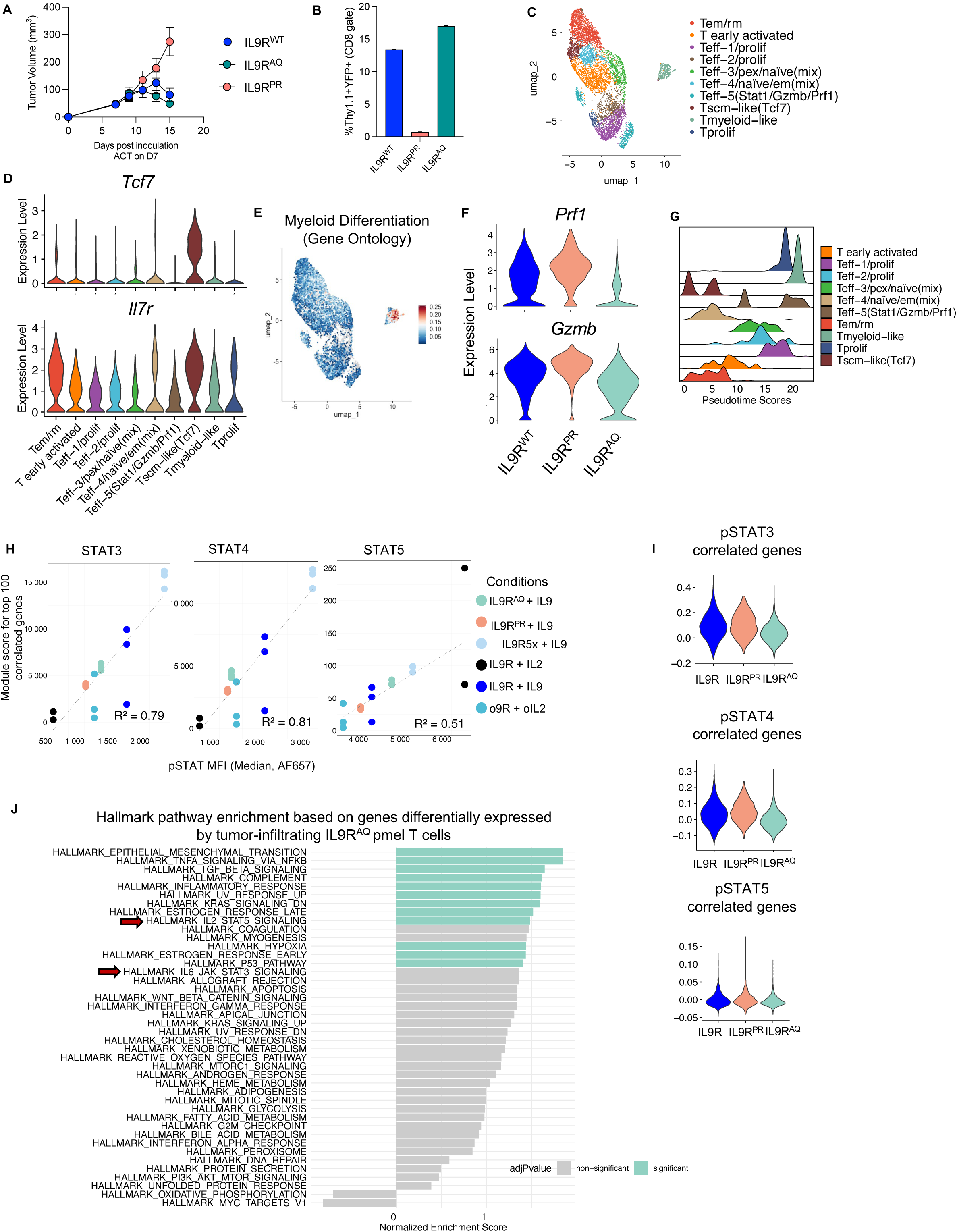
IL-9R variants reveal STAT1 as a rheostat of T cell differentiation, related to Figure 6. (A) Tumor growth (mean ± SEM) for the experiment described in (A). Data is representative of at least two independent experiments. (B) Transduced (Thy1.1^+^YFP^+^) pmel T cells as a percentage of total CD8 T cells in the tumor by treatment group for scRNAseq experiment from (A)-(B). (C) Merged UMAP plot based on scRNAseq of of n=6,706 cells pmel T cells transduced with IL-9R^WT^, IL-9R^AQ^, or IL-9R^PR^. Ten major clusters colored according to annotation. See related Figure 6A. (D) Violin plots summarizing single cell expression of stem and memory markers, *Tcf7* (top) and *Il7r* (bottom), by cluster. (E) Feature plot of module score defined by the Gene Ontology Myeloid Differentiation gene set demonstrating high expression exclusively in the Tmyeloid-like cluster. (F) Violin plots summarizing single cell expression of effector molecules, *Prf1* (top) and *Gzmb* (bottom), by treatment group. (G) Ridgeplot of pseudotime scores for single cell data from (A) organized by annotated cluster. (H) Scatterplot depicting the relationship between the pSTAT module scores (y-axis) and pSTAT phosphorylation levels (x-axis) in vitro. Module scores were calculated from the expression of 100 genes most strongly correlated with STAT phosphorylation in vitro. Shown are biological triplicates for the gene expression data, colored by sample condition. For STAT phosphorylation data, shown are the averages of technical duplicates. See also Figure 6K. (I) Projection of pSTAT module scores from (H) onto scRNA-seq data from Figure 6A. (J) Hallmark gene set enrichment analysis based on genes differentially expressed among tumor infiltrating IL-9R^AQ^ pmel T cells from Figure 6A.

**Figure S7.**
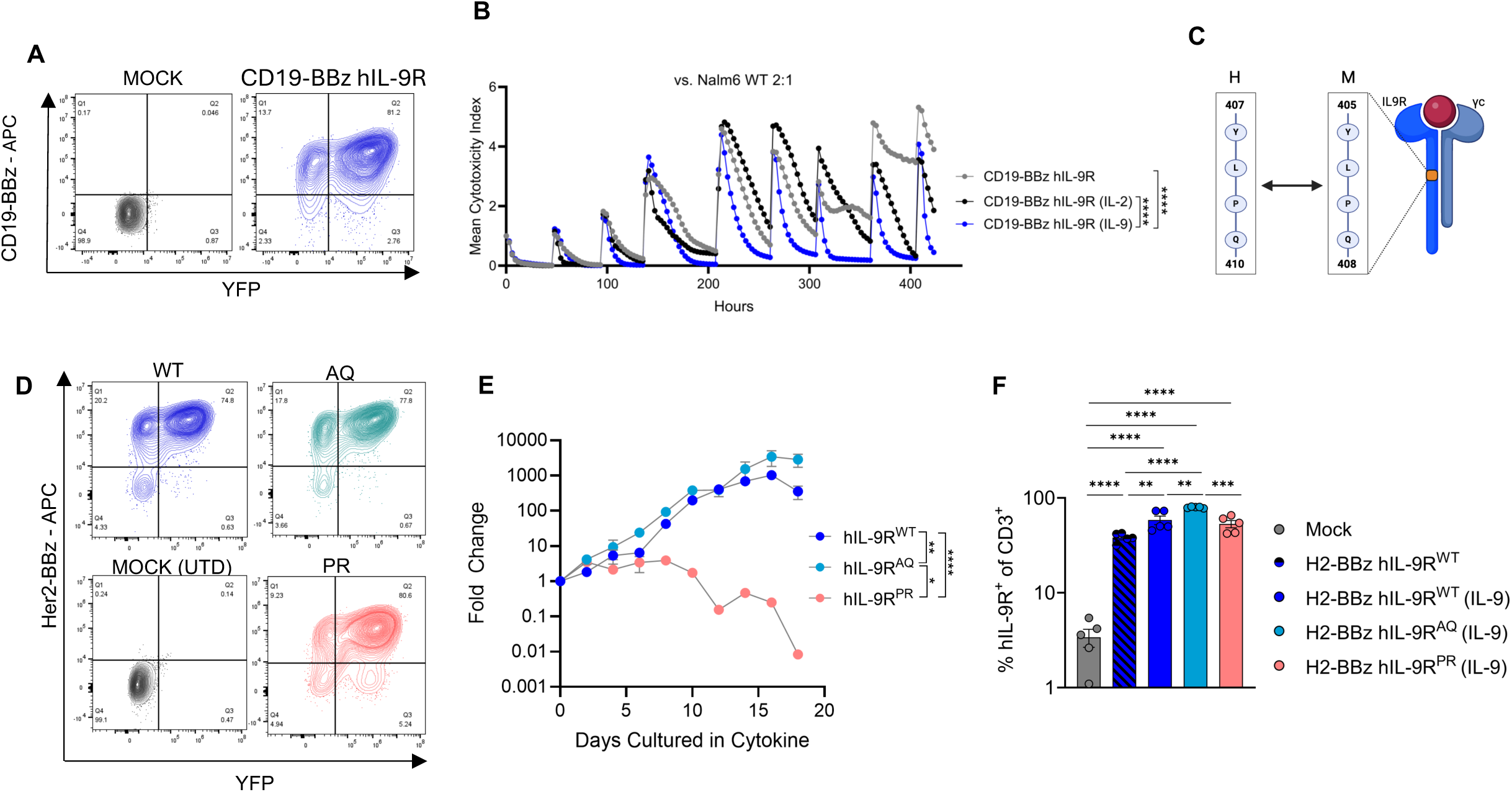
Phenotypic and functional impact of CAR T cells signaling through IL-9R and IL-9R variants, related to Figure 7. (A) Expression of second generation CD19 CAR (CD19-BBz) and human IL-9R after co-transduction as measured by idiotype staining and YFP expression, respectively. (B) In vitro cytotoxicity of second generation CD19 CAR (CD19-BBz) co-transduced with human IL-9R in a repetitive tumor challenge with NALM6 leukemia at a 2:1 E:T ratio. Cocultures were supplemented with IL-2, IL-9 or no cytokine. Shown is the mean cytotoxicity index as measured on the Incucyte SX5 based on tumor fluorescence (GFP). (C) Schematic demonstrating the conserved amino acids following the phosphotyrosine site of the intracellular domain of the human and mouse IL-9R. (D) Expression of second-generation Her2 CAR (Her2-BBz) and human IL-9R after co-transduction as measured by idiotype staining and YFP expression, respectively. (E) In vitro proliferation of human IL-9R transduced T cells grown in complete media supplemented with IL-9 (10nM). (F) Percentage of IL-9R^+^ (YFP^+^) CAR T cells as a percentage of total human T cells twelve days after tumor inoculation (8 days after ACT) from (K). See (J) *P < 0.05, **P < 0.005, ***P < 0.0005, ****P < 0.0001 (one-way ANOVA for B).

